# Tetravalent dengue mRNA vaccination orchestrates balanced four-serotype immunity with durable memory and protection in mice

**DOI:** 10.64898/2026.11.09.765416

**Authors:** Fahima Akther, Yago R. Dalben, John Villalpando, Kate S. Levine, Emma Buck, Jordan Polster, Zixi Yin, Hiromi Muramatsu, Sachchidanand Tiwari, Sowmya Sriram, Craig B. Wilen, Nikhil S. Joshi, Ryan P. McNamara, Daniela Weiskopf, Longping V. Tse, Norbert Pardi, David R. Martinez

**Affiliations:** Department of Immunobiology, Yale School of Medicine, New Haven, CT, USA; Department of Molecular Microbiology and Immunology, Saint Louis University School of Medicine, Saint Louis, Missouri, MO, USA; La Jolla Institute for Immunology, La Jolla, CA, USA; Department of Immunology and Infectious Diseases, Harvard T.H. Chan School of Public Health, Boston, MA, USA; Department of Microbial Pathogenesis, Yale School of Medicine, New Haven, CT, USA; Center for Infection and Immunity, Yale School of Medicine, New Haven, CT, USA; Department of Microbiology, University of Pennsylvania, Philadelphia, PA, USA; Department of Medicine, Division of Infectious Diseases and Global Public Health, University of California San Diego (UCSD), La Jolla, CA 92037, USA

**Keywords:** Dengue virus, Tetravalent vaccine, mRNA-LNP vaccine, Durable memory, Immune imprinting, Germinal center

## Abstract

An effective dengue vaccine must establish durable immunity across four dengue virus (DENV) serotypes, evade antibody-dependent enhancement (ADE), and overcome immune imprinting from prior exposure. Here, we developed Tet_prM-E_ mRNA-LNP, a tetravalent vaccine comprising LNP-formulated nucleoside-modified mRNAs encoding DENV1-4 precursor membrane-envelope (prM-E) antigens, and evaluated its immunogenicity, durability, protection and breath across serotype-specific immune backgrounds. Three immunization strategies elicited neutralizing titers greater than 1:1500 against each serotype. Passive transfer of immune sera conferred 100% survival and reduced viremia after lethal DENV2 challenge in immunocompromised mice. Vaccination induced DENV envelope (env)-specific CD8+T cells and CD4-dependent germinal-center responses. DENV env-binding antibodies persisted for 34 weeks alongside DENV env-specific memory B cells. Vaccine-elicited sera showed no measurable in vitro ADE. Although monovalent priming initially biased antibody responses, Tet_prM-E_ mRNA-LNP vaccination broadened neutralization across DENV1-4 regardless of priming serotype. Together, Tet_prM-E_ mRNA-LNP elicits broad, durable, and protective immunity despite serologic constraints from pre-existing immunity.

## Introduction

Dengue virus (DENV) is a major and expanding global health threat, causing an estimated 400 million infections annually and circulating across more than 130 countries, particularly in tropical and subtropical regions worldwide (*1*). DENV is a mosquito-borne, positive-sense single-stranded RNA virus transmitted primarily by *Aedes aegypti* and *Aedes albopictus* and comprises four antigenically distinct serotypes (DENV1-4) (*2*). Infection induces durable protection against the infecting serotype but provides incomplete and transient cross-protection against heterologous serotypes (*3*). Consequently, subsequent infection with a different DENV serotype can increase the risk of severe disease (*3*). A major mechanism implicated in this risk is thought to be antibody-dependent enhancement (ADE), in which pre-existing cross-reactive antibodies binding insufficiently neutralize a heterologous virus that can promote infection of Fcγ receptor-bearing cells, potentially increase viral replication and contribute to severe dengue disease (*4*). This complex interplay between protective and potentially disease-enhancing antibody responses presents a fundamental challenge for dengue vaccine development: immunity must be sufficiently broad, balanced, and durable to protect simultaneously against all four serotypes.

The DENV genome encodes three structural proteins: capsid (C), precursor membrane (prM), and envelope (env); and seven nonstructural proteins (NS1, NS2A, NS2B, NS3, NS4A, NS4B, and NS5) that coordinate viral replication, assembly, and interactions with host immunity (*2*). Among the DENV structural proteins, prM and env are particularly relevant to vaccine design. The prM protein associates with env during virion biogenesis and facilitates its proper folding and transport (*5*). Co-expression of prM and env is sufficient to support the formation and secretion of virus-like particles that display env in a virus-associated context (*6*). Env is a major antigenic target of neutralizing antibodies and therefore represents a central component of protective humoral immunity (*7, 8*). Together, these properties provide a strong rationale for selecting the prM-env (prM-E) region as the antigenic component of DENV vaccine constructs (*5, 7*). This principle is reflected in current tetravalent live-attenuated vaccines, although their strategies for presenting prM-E differ. CYD-TDV (Dengvaxia) expresses DENV1-4 prM-E in a yellow fever 17D backbone, whereas TAK-003 uses an attenuated DENV2 backbone together with chimeric viruses expressing prM-E from DENV1, DENV3, and DENV4 (*7, 9*). TV003/TV005 instead combines attenuated DENV1, DENV3, and DENV4 viruses with a DENV2/4 chimera expressing DENV2 prM-E (*9*). These vaccines have established the feasibility of targeting prM-E to generate protective immunity. However, achieving uniformly strong and durable protection across all four serotypes remains challenging (*7, 9*). Because tetravalent live-attenuated vaccines rely on replication of multiple viral components, differences in replication fitness, and immunodominance can influence the magnitude and balance of serotype-specific immune responses (*10*). Consequently, vaccine performance can vary across DENV serotypes and with pre-existing DENV immunity. Together, these observations support the prM-E region as a rational antigenic target for dengue vaccine development while highlighting the need for approaches that reduce dependence on differential viral replication and enable controlled presentation of antigens from all four serotypes, with the goal of promoting balanced and durable tetravalent immunity.

The global deployment of nucleoside-modified mRNA-lipid nanoparticle (mRNA-LNP) vaccines against corona virus disease (COVID)-19 established mRNA as a clinically viable vaccine platform and highlighted several features attractive for emerging and antigenically complex viral pathogens, including rapid antigen design, cell-free manufacturing, and the ability to combine multiple independently encoded antigens within a single formulation (*11–13*). These properties have motivated the development of mRNA-based vaccines against DENV, although progress remains largely preclinical. Early studies demonstrated that modified mRNA encoding conserved DENV1 nonstructural protein epitopes could elicit protective CD8+ T cell immunity in the absence of neutralizing antibodies, providing proof of concept for antibody-independent protection (*14*). Other approaches targeting DENV2 prM-E, E80, or NS1 demonstrated that mRNA vaccination can generate neutralizing antibodies and/or T cell responses and protect against homologous viral challenge (*15*). Similarly, a DENV1 prM-E mRNA vaccine elicited neutralizing antibodies and serotype-specific CD4^+^/CD8^+^ T cell responses (*7*). More recently, a multicomponent vaccine incorporating env domain III and NS1 antigens from multiple serotypes demonstrated the feasibility of generating tetravalent responses; however, neutralizing activity remained markedly lower against DENV2 and DENV3 than against DENV1 and DENV4, underscoring the challenge of achieving balanced immunity across serotypes (*16*). Structure-guided modification of DENV2 env protein, with a targeted N8R mutation to disrupt enhancing epitopes while preserving neutralizing domain, has also been explored to reduce enhancing antibody responses, although reductions in ADE were restricted to selected serum dilutions (*17*). Despite this progress, whether mRNA vaccination can generate balanced tetravalent immunity that remains durable as antibody responses wane remains unresolved. This question extends beyond peak neutralizing antibody titers to the establishment of germinal center (GC) and memory B cell (MBC) responses, cellular immunity, and the potential for ADE over time. Moreover, pre-existing serotype-specific immunity may reshape subsequent responses to heterologous DENV antigens, adding another challenge to maintaining tetravalent balance (*18–20*) These dimensions of immunity have yet to be comprehensively defined within a single tetravalent DENV mRNA vaccine platform.

To address these gaps, we developed a tetravalent DENV nucleoside-modified mRNA-LNP vaccine encoding the prM-E antigens of all four DENV serotypes and established a comprehensive preclinical framework to determine whether a single vaccine could achieve the breadth, balance, immune maturation, and durability that remain incompletely defined for DENV mRNA vaccination. We also define whether tetravalent DENV prM-E mRNA-LNP vaccination can establish balanced immunity that matures, persists, remains functionally protective, and is maintained in the context of pre-existing DENV immunity.

## Results

### Development of a tetravalent mRNA-LNP vaccine against DENV and evaluation of immunogenicity in a preclinical model

Prior to development of the tetravalent formulation, we performed antigen down-selection and monovalent immunogenicity studies, which identified prM-E as the lead immunogen and demonstrated serotype-specific antibody responses, as detailed in the supplementary results (**Figs. S1 and S2**). Based on these findings, we formulated the tetravalent DENV prM-E (Tet_prM-E_) mRNA-LNP vaccine by combining equal amounts of the four serotype-specific monovalent prM-E mRNA-LNPs into a single formulation **(Fig. 1A)**. Expression of prM and E proteins from the DENV1-4 prM-E mRNA constructs was confirmed by Western blot analysis (**Fig. S3A**). To evaluate the immunogenicity of Tet_prM-E_ mRNA-LNP, mice were immunized with 5 µg, comprising 1.25 µg of each serotype-specific prM-E mRNA-LNP, using a prime-boost-boost regimen at week 0, 4, and 8 (**Fig 1B**).

**Figure 1.**
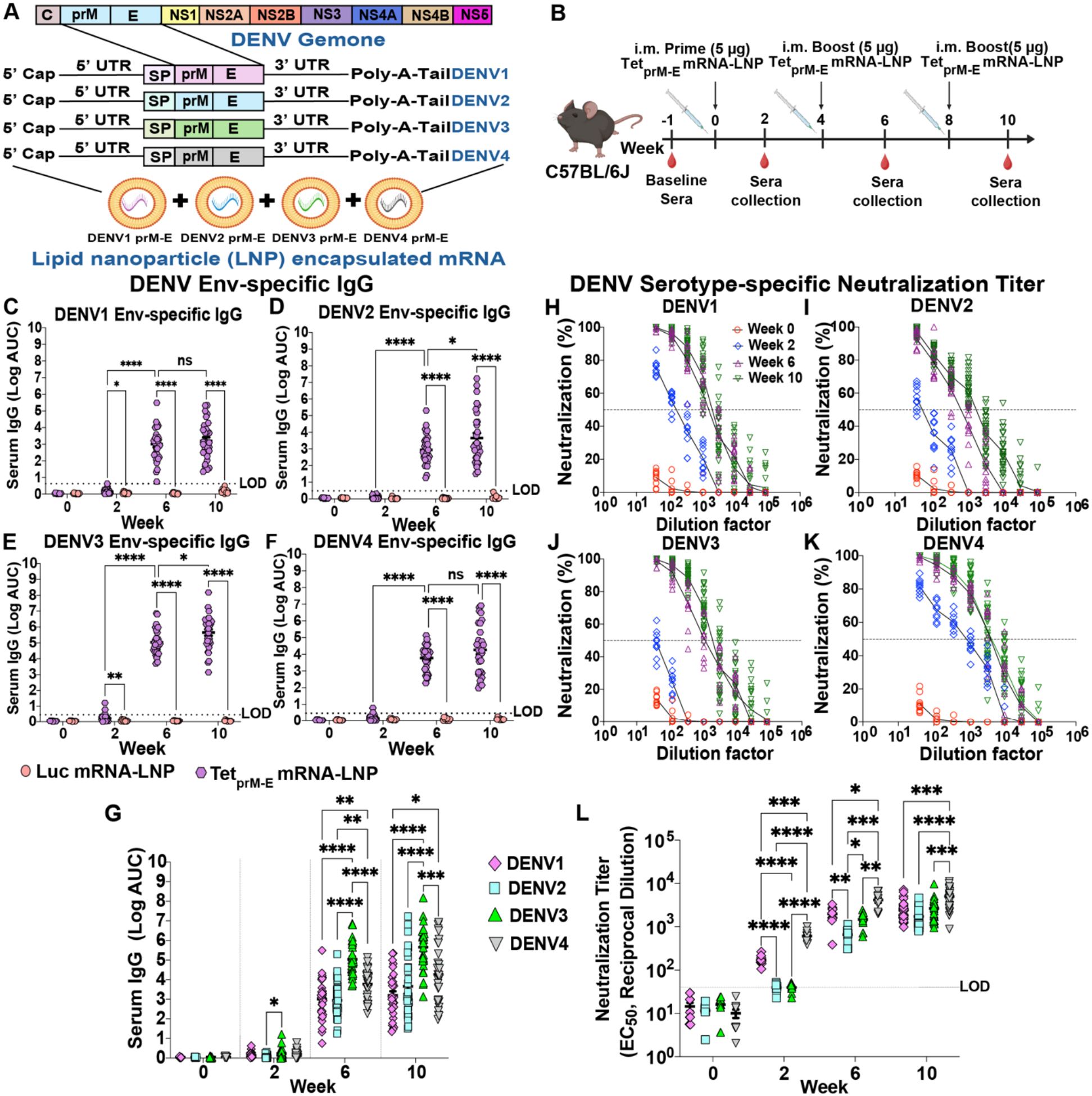
Tetravalent DENV prM-E (Tet_prM-E_) mRNA-LNP vaccination induces binding and neutralizing antibody responses against all four DENV serotypes. **(A)** Schematic of the DENV genome and Tet_prM-E_ mRNA-LNP (lipid nano particle) vaccine design. Individual mRNAs encoding the precursor membrane (prM) and envelope (env) proteins (PrM-E) of DENV1, DENV2, DENV3, or DENV4, preceded by a signal peptide (SP), were independently encapsulated in LNPs and combined to generate the tetravalent formulation. **(B)** Immunization and serum-collection schedule of Tet_prM-E_ mRNA-LNP vaccine. C57BL/6J mice were immunized intramuscularly (i.m.) with 5 μg of Tet_prM-E_ mRNA-LNP at weeks 0, 4, and 8. Serum was collected at baseline before priming and at weeks 2, 6, and 10, which was two weeks after each immunization. Serum IgG binding to the env proteins of DENV1 **(C)**, DENV2 **(D)**, DENV3 **(E)**, and DENV4 **(F)** was measured by antigen-specific ELISA at the indicated time points and expressed as log-transformed area under the curve (AUC). Mice receiving luciferase (luc) mRNA-LNP served as a negative control. The ELISA limit of detection (LOD) was defined as, LOD = mean of the blank + 3 × SD of the blank. **(G)** Comparison of env-specific IgG responses among DENV1-4 following tetravalent vaccination at the indicated time points. Neutralizing activity of serum collected at weeks 0, 2, 6, and 10 against DENV1 **(H)**, DENV2 **(I)**, DENV3 **(J)**, and DENV4 **(K)** infection. Neutralization is shown as the percentage of infection neutralized across serial serum dilutions; dashed lines indicate 50% neutralization. **(L)** Serotype-specific neutralizing antibody titers against DENV1-4 at the indicated time points, expressed as the reciprocal serum dilution producing 50% neutralization (EC_50_). Dashed lines indicate the assay LOD at a 1:40 dilution. The tetravalent vaccination study was performed across three independent cohorts (cohort 1, n = 10; cohort 2, n = 15; cohort 3, n = 5), for a combined total of 30 mice. Data are presented as mean ± SEM, with each symbol representing an individual mouse. Statistical significance was assessed using a mixed-effects model with the Geisser-Greenhouse correction for repeated measures, followed by Tukey’s multiple-comparisons test. \**P* < 0.05, \*\**P* < 0.01, \*\*\**P* < 0.001, \*\*\**P* < 0.0001; ns, not significant.

The tetravalent vaccination study was performed across three independent cohorts of mice (cohort 1, n = 10; cohort 2, n = 15; cohort 3, n = 5). This independent-cohort design allowed us to confirm that the immunogenicity of the Tet_prM-E_ mRNA-LNP vaccine was consistently reproduced across separate vaccination experiments. To determine whether the Tet_prM-E_ mRNA-LNP vaccine elicited antibody responses against all four DENV serotypes, we first evaluated the kinetics and breadth of serum IgG responses following vaccination. Sera were collected two weeks after the prime and after each subsequent boost and analyzed by antigen-specific ELISA using recombinant env proteins from DENV1-4. Recombinant proteins were selected to provide standardized antigen preparations across DENV serotypes and to minimize potential variability associated with whole-virion antigen preparations, which could confound comparisons of the immunogenicity of the tetravalent vaccines (*21, 22*). Luciferase (luc) mRNA-LNP immunized mice were served as negative control.

Following the first boost, a pronounced increase in IgG responses were observed, with mean IgG levels reaching 3.01 ± 0.19, 2.95 ± 0.17, 5.04 ± 0.15, and 3.78 ± 0.14 Log AUC (area under the curve) against DENV1, DENV2, DENV3, and DENV4 env, respectively **(Fig. 1C-F)**. Following the second boost, responses further sustained or increased to 3.20 ± 0.21, 3.66 ± 0.29, 5.65 ± 0.20, and 4.27 ± 0.27 Log AUC against DENV1-4 env, respectively. Luc mRNA-LNP control remained at or near background levels. Together, these results demonstrate that booster immunization with Tet_prM-E_ mRNA-LNP vaccine markedly enhances dengue env-binding antibody responses against each DENV serotype. Moreover, direct comparison across serotypes showed low and comparable responses after priming, followed by marked increases after boosting, with DENV3 exhibiting the highest binding response (**Fig. 1G**). Whole-virion ELISA similarly demonstrated IgG binding to all four DENV serotypes, with the highest binding observed for DENV2 (Fig. S3B).

We next determined whether vaccine-elicited antibodies neutralized all four DENV serotypes. Neutralization was assessed longitudinally against mature DENV1 (Vietnam/2008/Yale-DH144), DENV2 (Indonesia/1984/Yale-DH069), DENV3 (Cambodia/2011/Yale-DH166), and DENV4 (Venezuela/1990/Yale-DH127) using sera from vaccinated mice **(Fig. 1H-K)**. Viruses propagated in Vero-furin-overexpressing cells were used to enrich for mature virions with reduced uncleaved prM (*23, 24*). Minimal neutralizing activity was detected at baseline, whereas vaccination induced neutralizing antibodies that increase after each boost. Following the first boost, mean neutralization titers reached approximately 2,000 ± 260, 660 ± 79, 1,500 ± 210, and 4,100 ± 560 against DENV1-4, respectively; following the second boost, titers increased to 2,500 ± 270, 1,700 ± 160, 2,700 ± 320, and 5,000 ± 470, respectively (**Fig 1L**). DENV4 elicited the highest neutralization titer, whereas DENV1-3 were not significantly different after the second boost. Notably, although DENV3 showed the highest env-binding IgG response, this did not correspond to the highest neutralizing activity. However, with standard virus stocks grown in Vero-81 cells containing mature and partially immature virions, DENV3 showed the highest neutralization at peak immunogenicity (Fig. S3C, D). Together, these findings demonstrate that Tet_prM-E_ mRNA-LNP vaccination induces broad neutralizing antibody responses against all four DENV serotypes and that binding magnitude does not necessarily predict relative neutralization potency.

### CD4+ T cell help shapes germinal center responses to tetravalent DENV prM-E mRNA-LNP vaccination

Although vaccination can rapidly induce antigen-specific antibody responses, the establishment of GC is likely critical for the progressive selection and maturation of high-affinity B-cell clones and the development of durable humoral immunity (*25, 26*). GC formation is tightly dependent on coordinated interactions between antigen-activated B cells and CD4^+^ T cells, which provide essential signals for B-cell expansion, selection, and differentiation within the GC (*27*). Moreover, mRNA-LNP vaccination can induce durable GC responses (*28*), we examine whether Tet_prM-E_ mRNA-LNP-induced GC formation required CD4^+^ T cell help by depleting CD4^+^ T cell during booster immunization (**Fig. 2A)**. CD4^+^ T cell depletion was efficient in both vaccine-draining inguinal lymph nodes (dLN) **(Fig. 2B**) and spleen (**Fig 2G**). Representative flow cytometry plots showed near-complete loss of CD3^+^CD4^+^ T cells following an anti-CD4+ depleting antibody (GK1.5) treatment compared with isotype controls in both dLN and spleen, (**Fig 2B, G**, and **Fig. S4A)**. In dLN, CD4^+^ T cells decreased from 30-35% of live cells to <1% after depletion, corresponding to an approximately 99% reduction in CD4^+^ T-cell frequency **(Fig. 2C)**., with similarly profound reduction was observed in the spleen (**Fig. 2H**).

**Figure 2:**
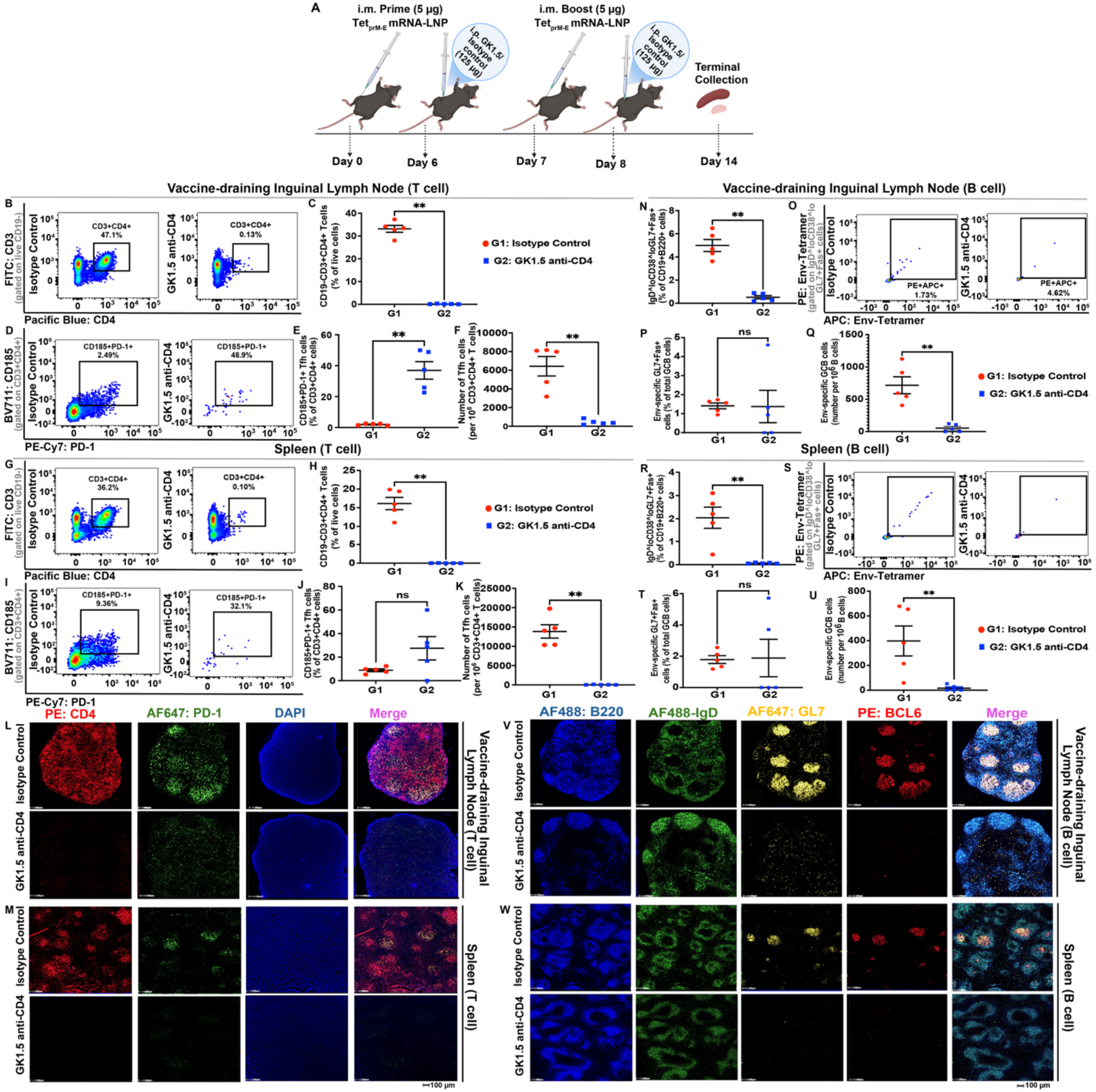
CD4^+^ T cell depletion impairs T follicular helper cells (Tfh) and germinal center (GC) B cell responses following tetravalent DENV prM-E (Tet_prM-E_) mRNA-LNP vaccination. **(A)** Experimental design of CD4+ T cell depletion. Mice were primed intramuscularly (i.m.) with 5 μg Tet_prM-E_ mRNA-LNP on day 0 and boosted with the same dose on day 7. Mice received either isotype control antibody (G1; 125 μg per dose) or GK1.5 anti-CD4 depleting antibody (G2; 125 μg per dose) on days 6 and 8. Vaccine-draining inguinal lymph nodes and spleens were collected on day 14 for analysis. Flow-cytometric analysis of T cell responses in vaccine-draining inguinal lymph nodes: the representative plots show CD3^+^CD4^+^ T cells in both groups **(B)**, quantification of the frequency of CD3^+^CD4^+^ T cells among live CD19⁻ cells **(C)**, representative plot of CD185^+^PD-1^+^ Tfh cells among CD3^+^CD4^+^ T cells in both groups **(D)**, the frequency of CD185^+^PD-1^+^ Tfh cells among CD3^+^CD4^+^ T cells **(E)**, and the number of Tfh cells per 1 × 10^6^ CD3^+^CD4^+^ T cells **(F)**. Flow-cytometric analysis of splenic T-cell responses: the representative flow-cytometry plots of CD3^+^CD4^+^ T cells in both groups **(G)**, quantification of CD3^+^CD4^+^ T cell frequency **(H)**, representative plot of CD185^+^PD-1^+^ Tfh cells **(I)**, CD185^+^PD-1^+^ Tfh cell frequency **(J)**, and Tfh cell number per 1 × 10^6^ CD3^+^CD4^+^ T cells **(K)**. Representative immunofluorescence images of vaccine-draining inguinal lymph nodes **(L)** and spleens **(M)** from isotype control- and GK1.5 anti-CD4-treated mice, stained for CD4 (red), PD-1 (green), and DAPI (blue) with merged images shown. Analysis of GC B-cell responses in vaccine-draining inguinal lymph nodes: frequency of IgD^loCD38^loGL7^+^Fas^+^ GC B cells among CD3-CD19^+^B220^+^ cells **(N)**, representative flow-cytometry plots of DENV3 env-specific GC B cells **(O)**, frequency of env-specific GL7^+^Fas^+^ cells among total GC B cells **(P)** and number of env-specific GC B cells per 1 × 10^6^ B cells **(Q)**. Analysis of GC B cell responses in splenic B cells: frequency of IgD^loCD38^loGL7^+^Fas^+^ GC B cells among CD3-CD19^+^B220^+^ cells **(R)**, representative DENV3 env-specific GC B cells **(S)**, the frequency of env-specific GL7^+^Fas^+^ cells among total GC B cells **(T)**, and the number of env-specific GC B cells per 1 × 10^6^ B cells **(U)**. Representative immunofluorescence images of vaccine-draining inguinal lymph nodes **(V)** and spleens **(W)** stained for B220 (blue), IgD (green), GL7 (yellow), and BCL6 (red), with merged images shown. Each symbol represents an individual mouse; error bars indicate mean ± SEM. Statistical comparisons between groups were performed using a two-tailed Mann-Whitney U test. ns, not significant; \*\**P* < 0.01. Scale bar = 100 µm.

We next examined the residual CD4^+^ T cell compartment for a T follicular helper (Tfh)-associated CD185+PD-1+ phenotype. Although the relative frequency of CD185^+^PD-1^+^ cells among residual CD4^+^ T cells were increased in dLN after depletion (**Fig. 2D, E**), absolute Tfh abundance was markedly reduced, from 6,431 ± 1,040 to 432 ± 117 cells (**Fig. 2F**) per 1 × 10^6^ CD3+CD4+ T cells. A similar pattern was observed in the spleen, where Tfh frequency within the residual CD4^+^ T cell compartment was unchanged, but absolute numbers decreased from 13,838 ± 1,731 to 47 ± 25 cells (**Fig. 2I-K**). These results corresponded to approximately 93% and 99% reductions in Tfh abundance in dLN and spleen, respectively. Immunofluorescence confirmed the marked loss of CD4^+^ T cells and associated PD-1 staining in both tissues following depletion (**Fig. 2L, M**).

Further, we assessed the impact of CD4+ T cell depletion on GC B cell formation. In dLN, total GC B cells (IgD^loCD38^loGL7+Fas+) was markedly reduced following CD4^+^ T cell depletion (**Fig. 2N**; gating strategy in **Fig. S4B**). Although the frequency of DENV3 env-specific cells within the residual GC B cell compartment was unchanged (**Fig. 2O, P**), their absolute number was profoundly reduced from 717 ± 131 to 55 ± 26 cells per 1 × 10^6^ B cells after depletion (**Fig. 2Q**), indicating a substantial loss of the antigen-specific GC B cell population. A comparable pattern was observed in the spleen, where total GC B cell frequencies were reduced following CD4^+^ T cell depletion (**Fig. 2R**). The proportion of DENV3 env-specific cells within residual GC B cells was unchanged (**Fig. 2S, T**), whereas their absolute number decreased from 398 ± 121 to 16 ± 11 cells per 1 × 10^6^ B cells (**Fig. 2U**). Immunofluorescence further demonstrated loss of GL7^+^BCL6^+^ GC structures in both dLN and spleen following CD4^+^ T cell depletion (**Fig. 2V, W**). Together, these findings demonstrate that CD4^+^ T cell help is required to sustain vaccine-induced total and DENV3 env-specific GC B cell responses across lymphoid tissues.

### Vaccine-elicited antibodies confer protection against lethal DENV2 challenge in vivo and show limited ADE in vitro

To determine whether vaccine-elicited antibodies confer protection against DENV2 infection in vivo, six-week-old AG129 mice were passively immunized with Tet_prM-E_ mRNA-LNP vaccine-derived immune sera prior to challenge with a lethal dose of 1 × 10^6^ FFU of mouse-adapted DENV2 S221 per mouse **(Fig. 3A)**. Mice were monitored daily for changes in body weight and clinical signs of disease and were humanely euthanized upon reaching the predefined humane endpoint of 80% of their initial body weight or or the development of severe clinical symptoms. Following lethal DENV2 challenge, PBS-treated mice exhibited progressive body-weight loss. By 4 dpi, the weight loss had become pronounced and continued until animals reached the predefined weight-loss threshold or developed severe clinical signs, at which point they were humanely euthanized **(Fig. 3B)**. In contrast, mice that received the Tet_prM-E_ mRNA-LNP vaccine-derived antibodies-maintained body weights near baseline throughout the observation period, with only minor fluctuations. A similar protective weight profile was observed in mice treated with the EDE1-C10 dengue broadly neutralizing monoclonal antibody used as a positive control. Consistent with the progressive weight loss, PBS-treated mice began to succumb to infection at 5 dpi, and all animals reached the predefined humane endpoint by 7 dpi (**Fig. 3C)**. In contrast, mice that received either vaccine-derived antibodies or EDE1-C10 exhibited 100% survival throughout the 10-day observation period, demonstrating that passive transfer of vaccine-elicited antibodies protected against lethal DENV2 challenge.

**Figure 3.**
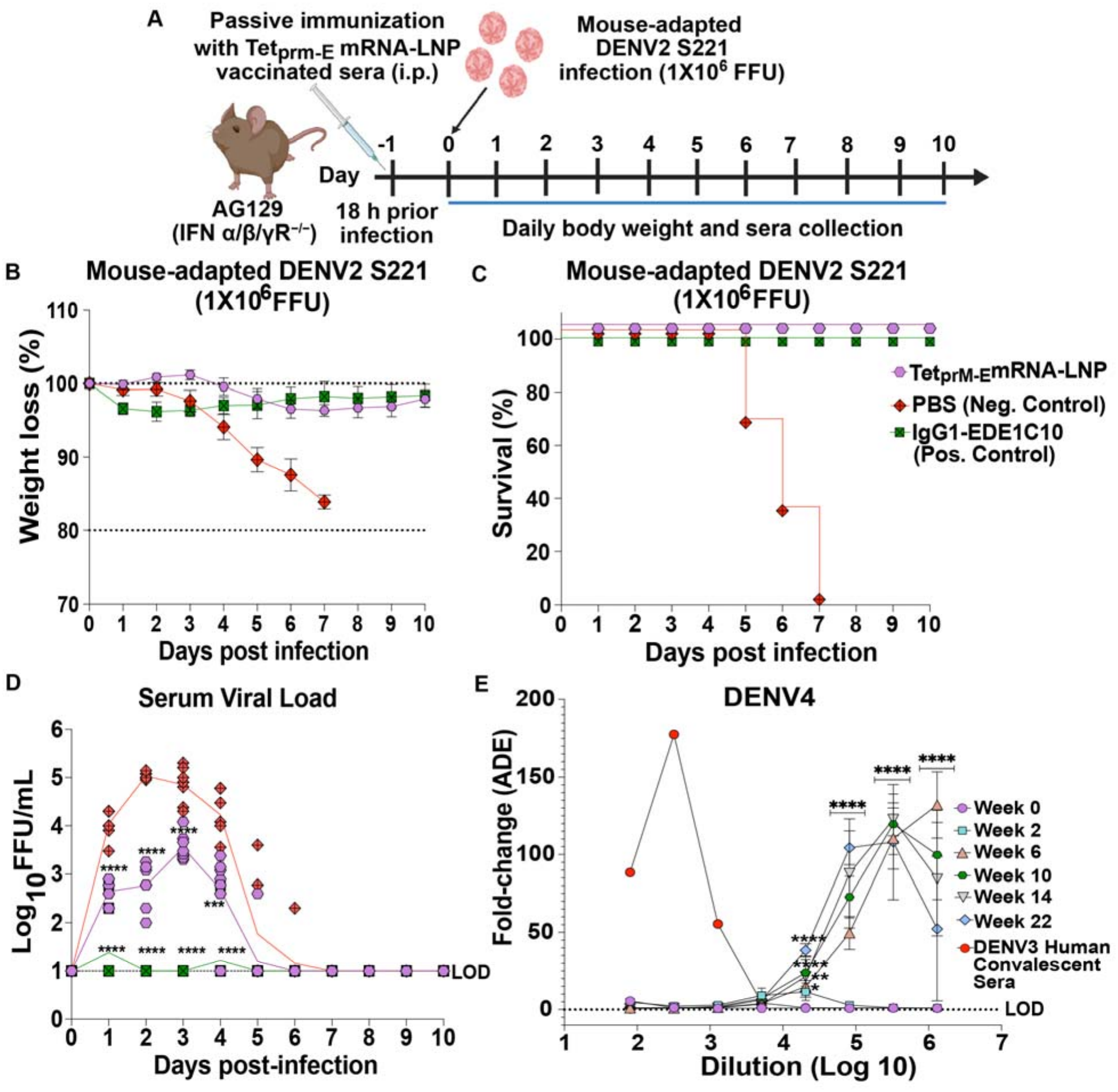
Antibodies elicited by tetravalent DENV prM-E (Tet_prM-E_) mRNA-LNP vaccination confer protection against lethal DENV2 challenge. **(A)** Schematic of the passive-transfer and DENV2 challenge study. Six-week-old male and female AG129 (IFN-α/β/γR^−/−^) mice received sera from Tet_prM-E_ mRNA-LNP-vaccinated mice via intraperitoneal (i.p.) injection 18 h before infection with 1 × 10^6^ focus-forming units (FFU) of mouse-adapted DENV2 S221. 1X PBS-treated mice served as negative controls, and the DENV-neutralizing monoclonal antibody IgG1-EDE1-C10 served as a positive control. Mice were monitored daily for body weight and survival, and sera were collected for measurement of viral load. **(B)** Body weight following DENV2 challenge, expressed as the percentage of initial body weight. **(C)** Kaplan-Meier survival curves following DENV2 challenge. **(D)** Serum viral loads following challenge, expressed as log10 FFU/mL. The assay limit of detection (LOD) is indicated by the dashed line. For the challenge study, Tet_prM-E_ mRNA-LNP and PBS groups contained n = 8 mice per group, and the IgG1-EDE1-C10 group contained n = 4 mice. Longitudinal data from the challenge study were analyzed using a mixed-effects model (REML) with the Geisser-Greenhouse correction, followed by Dunnett’s multiple-comparisons test. \*\*\**P* < 0.001, \*\*\*\**P* < 0.0001. **(E)** Antibody-dependent enhancement (ADE) activity of sera collected from Tet_prM-E_ mRNA-LNP-vaccinated mice at weeks 0, 2, 6, 10, 14, and 22 against DENV4. ADE was measured in K562 cells and is expressed as fold change in luminescence relative to the no-antibody control (cells and virus only) across serial serum dilutions. DENV3 human convalescent serum was included as a positive control. Data from the ADE study were analyzed using Two-way Anova, followed by Dunnett’s multiple comparison test. \**P* < 0.05, \*\**P* < 0.01, \*\*\*\**P* < 0.0001. All data are presented as mean ± SEM. Each symbol represents an individual mouse where applicable.

Protection from disease was accompanied by a significant reduction in circulating viral load **(Fig. 3D)**. In PBS-treated mice, viral titers increased rapidly following challenge and peaked at approximately 5 log_10_ FFU/mL at 2 dpi. In comparison, mice that received Tet_prM-E_ mRNA-LNP vaccine-elicited antibodies exhibited a substantially lower viral burden throughout the experiment, with a mean viral titer of approximately 2.9 log10 FFU/mL, representing an approximately 2.1 log_10_ reduction relative to PBS-treated animals. Viral titers in the vaccine-antibody group remained consistently lower than those observed in PBS-treated animals, with an approximately 1.5 log_10_ reduction. Viral titers then declined rapidly after 4 dpi, approaching the assay limit of detection by approximately 6 dpi. EDE1-C10 treated mice also had suppressed viral replication, with serum viral titers remaining close to the limit of detection throughout the acute phase of infection. Thus, passive transfer of Tet_prM-E_ mRNA-LNP vaccine-elicited antibodies significantly reduce viremia and protects against progressive weight loss and mortality.

As antibody-dependent enhancement (ADE) is a concern for investigational dengue virus vaccines, we measured whether Tet_prM-E_ mRNA-LNP vaccination elicited ADE antibodies. Human DENV3 convalescent serum, included as a heterotypic ADE control, showed pronounced enhancement of DENV4 infection at 1:320 dilutions. This is consistent with the well-described ability of heterotypic DENV antibodies at sub-neutralizing concentrations to enhance infection of FcγR-expressing cells (*29*). In contrast, sera collected from mice following tetravalent vaccination showed minimal enhancement at serum dilutions where convalescent DENV3 sera enhanced infection, with responses remaining close to the week 0 baseline through approximately the 1:10,000 dilution range. At dilution above 10,000, increased ADE was observed in post-vaccination sera, particularly at weeks 6, 10, 14, and 22, with peak enhancement reaching approximately 100-130-fold (**Fig. 3E**). Thus, vaccine-elicited sera showed minimal ADE across a broad dilution range, with enhancement becoming apparent only at the highest serum dilutions tested. Importantly, the dilutions at which in vitro ADE was observed with Tet_prM-E_ mRNA-LNP vaccine are unlikely to be representative of physiological relevant dilutions in humans where dengue serum titers of 1:21 to 1:80 are associated with ADE (*30*). This high-dilution enhancement is consistent with the concentration-dependent nature of in vitro ADE, in which progressive serum dilution eventually reduces antibody concentrations into a sub-neutralizing range that permits FcγR-mediated enhancement of infection (*29*).

### Tetravalent DENV prM-E mRNA-LNP vaccination elicits antigen-specific and functional T cell responses

In addition to antibody-mediated immunity, cellular immune responses may contribute to protection against DENV infection through recognition and elimination of virus-infected cells. We therefore investigated whether immunization with the Tet_prM-E_ mRNA-LNP vaccine elicited antigen-specific T cell responses. T cell responses were assessed by intracellular cytokine staining following 5 hour ex-vivo stimulation with peptide pools corresponding to DENV prM, E, or NS1 a negative control DENV antigen. Cytokine production was evaluated within CD4+ and CD8+ T cell populations by measuring IFN-γ, TNF-α, and IL-2. Luc mRNA-LNP-immunized animals served as controls, and the complete flow-cytometric gating strategy is provided in **Fig. S5**.

Antigen-specific cytokine production within the CD4+ T cell compartment was limited. Granzyme B (GrzB) expression was also assessed but showed no antigen-specific increase above background (**Fig 4A**). Frequencies of TNF-α+, IL-2+, IFN-γ+, and IFN-γ+TNF-α+ CD4+ T cells did not differ significantly between the Tet_prM-E_ mRNA vaccine and luc mRNA-LNP control groups following stimulation with prM, E, or NS1 peptide pools (**Fig. 4B-E)**. These data indicate modest vaccine-induced cytokine responses within the CD4^+^ T-cell compartment under the conditions examined.

**Figure 4.**
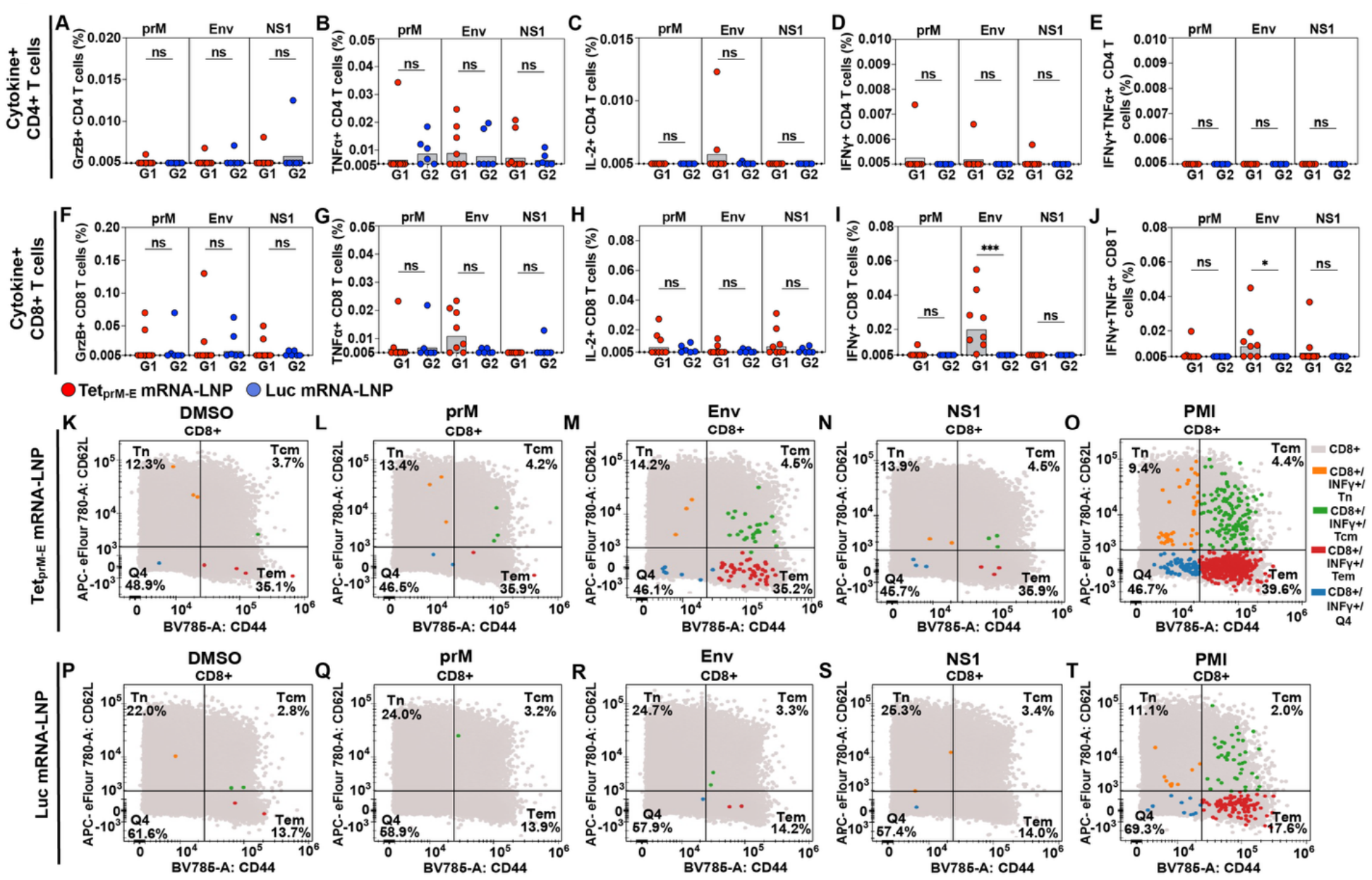
Tetravalent DENV prM-E (Tet_prM-E_) mRNA-LNP vaccination induces DENV antigen-specific T cell responses. Antigen-specific cytokine responses in CD4+ and CD8+ T cells following ex vivo stimulation in splenocytes with DENV prM, Env, or NS1 antigens. Splenocytes from mice immunized with Tet_prM-E_ mRNA-LNP (G1; *n* = 8) or luciferase (luc) mRNA-LNP (G2; *n* = 6) were stimulated with the indicated DENV antigens, and cytokine-producing T cells were quantified by flow cytometry. Frequencies of GrzB+ **(A)**, TNFα+ **(B)**, IL-2+ **(C)**, IFNγ+ **(D)**, and IFNγ+TNFα+ **(E)** CD4+ T cells. Frequencies of GrzB+ **(F)**, TNFα+ **(G)**, IL-2+ **(H)**, IFNγ+ **(I)**, and IFNγ+TNFα+ **(J)** CD8+ T cells. Data are presented as mean ± SEM, with each symbol representing an individual mouse. Statistical significance between G1 and G2 was assessed using a two-tailed Mann-Whitney U test. Representative flow-cytometry plots showing the differentiation phenotype of IFNγ+ CD8+ T cells following stimulation with DMSO negative control **(K, P)**, prM **(L, Q)**, Env **(M, R)**, NS1 **(N, S)**, and PMA/ionomycin (PMI) positive control **(O, T)** from Tet_prM-E_ mRNA-LNP-vaccinated **(K-O)** and luc mRNA-LNP control **(P-T)** mice. IFNγ+ CD8+ T cells were characterized based on CD44 and CD62L expression as naïve (Tn; CD44−CD62L+), central memory (Tcm; CD44+CD62L+), effector memory (Tem; CD44+CD62L−), or CD44−CD62L− (Q4) populations. Colored events indicate IFNγ+ cells within each phenotypic population. \**P* < 0.05, \*\**P* < 0.001; ns, not significant.

In contrast, vaccination elicited a pronounced antigen-specific CD8^+^ T cell response predominantly directed against DENV env. GrzB expression in CD8^+^ T cells likewise showed no antigen-specific increase above background (**Fig. 4F**). Following env peptide stimulation, Tet_prM-E_ mRNA-LNP-immunized animals exhibited significantly higher frequencies of IFN-γ^+^ CD8^+^ T cells than luc mRNA-LNP controls (**Fig. 4I**), whereas responses to prM or NS1 remained at or near background levels. TNF-α^+^ and IL-2^+^ CD8^+^ T-cell frequencies did not differ significantly between groups following stimulation with any peptide pool (**Fig. 4G, H**). Env-specific polyfunctional IFN-γ^+^TNF-α^+^ CD8^+^ T-cell responses were also significantly increased following Tet_prM-E_ mRNA-LNP vaccination, whereas responses to prM and NS1 did not differ between groups (**Fig. 4J**).

We next examined the memory phenotype of antigen-responsive CD8+ T cells based on CD44 and CD62L expression. The DMSO negative control yielded minimal background IFN-γ responses (**Fig 4K, P**), whereas the positive stimulation control (PMI; PMA/ionomycin) induced robust IFN-γ production (**Fig 4O, T**), confirming assay responsiveness. Following env peptide stimulation, IFN-γ+ CD8+ T cells were more evident in the Tet_prM-E_ mRNA-immunized animals than in controls (**Fig. 4M, R**) and were primarily localized within the effector-memory (T_EM; CD44+CD62L−) and central-memory (T_CM; CD44+CD62L+) compartments. Few IFN-γ+ CD8+ T cells were detected following prM or NS1 stimulation in Tet_prM-E_ mRNA-LNP vaccination in either group (**Fig. 4L, N, Q, S).** Collectively, these findings demonstrate that Tet_prM-E_ mRNA-LNP vaccination induces a detectable cellular immune response characterized predominantly by env-specific CD8+ T cells, including IFN-γ-producing and polyfunctional IFN-γ+TNF-α+ populations. The absence of a corresponding NS1-specific response further supports the antigen specificity of the vaccine-induced CD8+ T-cell response.

### Tetravalent DENV prM-E mRNA-LNP vaccination induces broad, durable humoral immunity and immune memory

To assess the durability of vaccine-induced humoral immunity and immune memory, mice immunized with Tet_prM-E_ mRNA-LNP were evaluated longitudinally through week 34 (**Fig. 5A**). Serum and spleens were collected at weeks 10, 14, 22, and 34 to assess the persistence of DENV-specific antibody and MBC responses.

**Figure 5.**
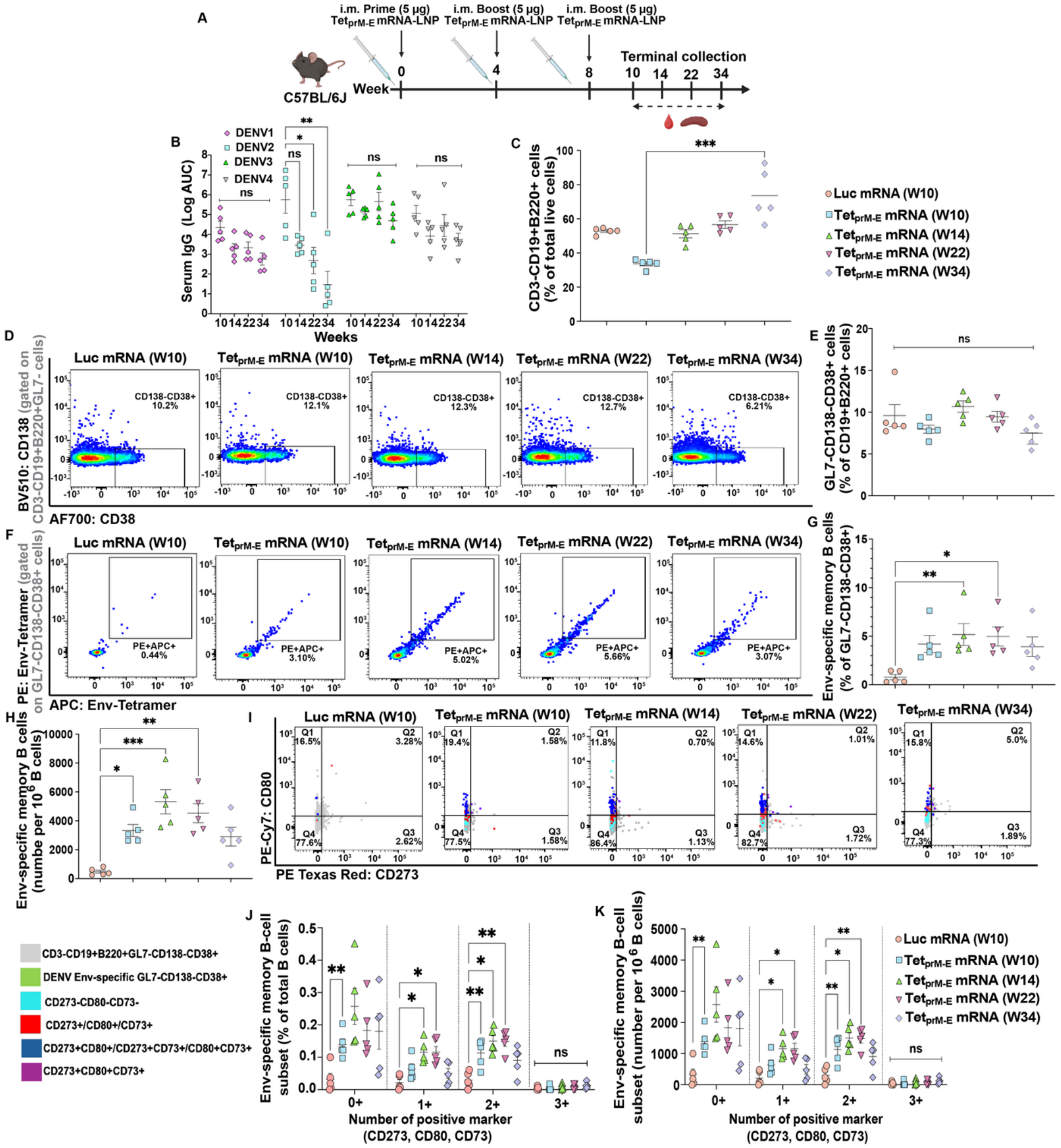
Durability of antibody responses and DENV-specific memory B cell (MBC) responses following tetravalent DENV prM-E (Tet_prM-E_) mRNA-LNP vaccination. **(A)** Schematic of the durability study. C57BL/6J mice were immunized intramuscularly (i.m.) with 5 μg Tet_prM-E_ mRNA-LNP at weeks 0, 4, and 8. Terminal sample collections were performed at weeks 10, 14, 22, and 34. **(B)** Longitudinal serum IgG binding responses against DENV-4 envelope (env) antigens at weeks 10, 14, 22, and 34, presented as log area under the curve (AUC). **(C)** Frequencies of CD3-CD19+B220+ cells among total live cells in luciferase (luc) mRNA-LNP control mice at week 10 and Tet_prM-E_ mRNA-LNP-vaccinated mice at weeks 10, 14, 22, and 34. **(D)** Representative flow cytometry plots showing identification of GL7-CD138-CD38+ MBCs within the CD3-CD19+B220+ B-cell population. **(E)** Frequencies of GL7-CD138-CD38+ memory B cells among CD3-CD19+B220+ cells. **(F)** Representative flow cytometry plots showing identification of DENV3 env-specific MBCs. **(G)** Frequencies of env-specific MBCs among GL7-CD138-CD38+ cells. **(H)** Absolute numbers of env-specific MBCs per 1 × 10^6^ B cells. **(I)** Representative flow cytometry plots showing CD273, CD80, and CD73 expression on env-specific MBCs. Distribution of env-specific MBC subsets according to the number of positive memory markers (CD273, CD80, and CD73), expressed as **(J)** percentage of total B cells and **(K)** number per 1 × 10^6^ B cells. Data are presented as mean ± SEM; *n* = 5 mice per group. Comparisons among multiple groups were performed using the Kruskal-Wallis test followed by Dunn’s multiple-comparisons test. When two factors were analyzed with repeated measurements, statistical significance was assessed using a mixed-effects model (REML) with the Geisser–Greenhouse correction, followed by Tukey’s multiple-comparisons test. ns, not significant; \**P* < 0.05, \*\**P* < 0.01, \*\*\**P* < 0.001.

DENV env-specific IgG responses were largely maintained over time, although the durability of binding antibodies differed among serotypes. DENV1-, DENV3-, and DENV4-specific IgG levels remained relatively stable from weeks 10 through 34, whereas DENV2-specific IgG responses declined at later time points **(Fig 5B)**. Despite this decline, binding IgG remained detectable against all four DENV serotypes throughout the follow-up period.

To determine whether the sustained antibody responses were accompanied by durable MBCs, splenic B cell populations were characterized longitudinally following Tet_prM-E_ mRNA-LNP vaccination. Total B cell frequencies and the broader GL7⁻CD138⁻CD38^+^ memory-like B-cell compartment remained largely stable over time (**Fig. 5C-E**; gating strategy in Fig. S6). We next assessed antigen-specific MBCs using a dual-labeled DENV3 env-tetramer probe consisting of biotinylated DENV3 env-dimers complexed with PE- and APC-conjugated streptavidin (**Fig. 5F**). DENV3 env-specific MBC frequencies were increased following vaccination and remained detectable from weeks 10 through 34 (**Fig. 5G**). Absolute enumeration similarly demonstrated persistence of DENV3 env-specific memory B cells through week 34, despite a modest decline at the latest time point (**Fig. 5H**).

To further characterize the memory response, DENV3 env-specific MBCs were evaluated for CD73, CD80, and CD273 (PD-L2) expression (**Fig. 5I**). Cells expressing one or two memory-associated markers were enriched following vaccination and persisted across multiple post-vaccination time points (**Fig. 5J, K**), whereas triple-positive cells remained infrequent. Collectively, these data demonstrate that Tet_prM-E_ mRNA-LNP vaccination induces durable serotype-spanning antibody responses and persistent DENV3 env-specific MBC populations.

### Systems serology defines the evolution and durability of the vaccine-induced antibody response

Further defining the quality and durability of the vaccine-induced humoral response, we performed longitudinal systems serology profiling. While antigen-specific antibody titers provide an important measure of response magnitude, they do not fully capture the breadth and qualitative features of the polyclonal antibody response (*31*). This approach was used to characterize antigen-specific antibody isotypes, subclasses, and Fc receptor-binding properties following Tet_prM-E_ mRNA-LNP vaccination.

Longitudinal profiling was performed at week 0, 2, 6, 10, 14, and 22, spanning baseline, post-prime, post-boost, peak immunogenicity, and later durability time points. Antibody responses against a panel of DENV env-derived antigens were characterized across multiple isotypes and subclasses, including Total IgG, IgG subclasses (IgG1, IgG2a, IgG2b, IgG2c, IgG3), IgA, and IgM, together with Fc receptor-binding features.

Z-score heat-map analysis showed low DENV-specific responses at baseline, followed by progressive expansion after the prime and booster immunizations (**Fig. 6**). Total IgG responses were detected across multiple DENV env-derived antigens, with particularly strong and sustained responses against DENV1 and DENV4 E-dimers (Fig. S8A). Vaccination also elicited distinct isotype kinetics, with an earlier IgM response and relatively stable IgA responses over time (**Fig. 6**). Responses to unrelated control antigens remained low, supporting antigen specificity. Broad DENV-specific responses persisted through weeks 14 and 22, and fold-change analysis relative to baseline similarly demonstrated sustained increases in DENV-specific IgG (Fig. S8B). In contrast, antibody responses against the unrelated control antigens HCMV gB, human serum albumin, and Ebola gP remained low, supporting the antigen specificity of the vaccine-induced response. The stratification of the longitudinal systems serology profiles by sex revealed comparable overall response trajectories in male and female mice, arguing against a sex-based influence on the immune outputs **(Fig S8 and S9)**.

**Figure 6.**
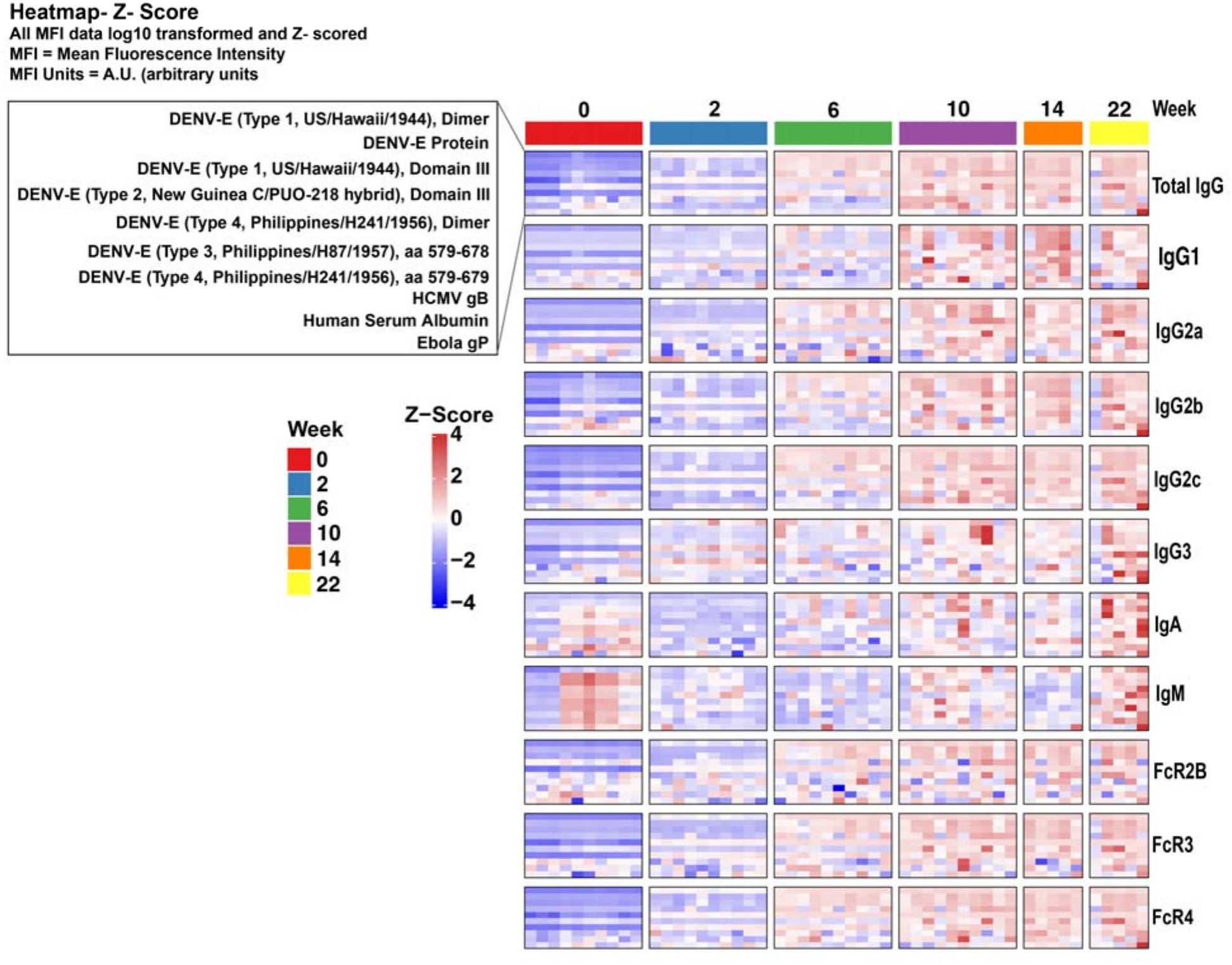
Longitudinal systems serology profiling of DENV-specific antibody responses. Serum antibody responses to DENV envelope (env)-derived antigens were characterized longitudinally using a multiplexed bead-based systems serology assay. Responses were measured at weeks 0, 2, 6, 10, 14, and 22 against the indicated DENV env antigens and control proteins, including HCMV gB, human serum albumin, and Ebola glycoprotein (gP). Antibody features included Total IgG, IgG subclasses (IgG1, IgG2a, IgG2b, IgG2c, and IgG3), IgA, IgM, and binding to FcγR2b, FcγR3, and FcγR4. Heatmap values represent Z-score-normalized antibody responses, with red indicating higher and blue indicating lower relative responses. Within each antibody feature, rows represent the indicated antigens and columns represent individual mice, grouped by collection time point as indicated by the colored bars above the heatmap. Weeks 0, 2, 6, and 10 include 10 mice per time point, whereas weeks 14 and 22 include 5 mice per time point.

Beyond antibody magnitude, longitudinal systems serology revealed a broader qualitative antibody profile encompassing multiple IgG subclasses and isotypes together with Fc receptor-binding features. FcγR2B-, FcγR3-, and FcγR4-binding responses developed following immunization and remained evident at later time points **(Fig 6)**. Composite humoral profiles of the vaccinated mice were then projected onto a principal component analysis (PCA) scatter plot. This unsupervised, multivariate analysis showed that most differences were along principal component axis 1 (PC1), which accounted for 58% of the total variance (**Fig S10A**). Notably, separations were based on immunization alone, with time since vaccination groups showing overlapping confidence intervals, but entirely separated from the baseline collection (week 0, gray circles). A biplot was created from the PCA to identify what features were driving the separation, particularly along PC1. The top 8 features were shown, which included humoral features directed against DENV env Type 1, 3, and 4, particularly Domain III of the env, or aa 579-679. Humoral features included IgG subclasses and FcγR-binding antibodies, further highlighting that Tet_prM-E_ mRNA-LNP vaccination was stimulating a binding and FcγR-binding humoral response (**Fig S10B**).

Collectively, these data demonstrate that Tet_prM-E_ mRNA-LNP vaccination induces a broad and multidimensional DENV-specific antibody response that evolves following sequential immunization and polyfunctionality remains durable for at least three months beyond peak immunogenicity, encompassing both antigen-specific antibody recognition and Fc receptor-binding features.

### Pre-Existing DENV immunity does not constrain tetravalent vaccine responses

Pre-existing immunity is a critical consideration in dengue vaccination because individuals in endemic regions are frequently exposed to one or more DENV serotypes before vaccination. Such prior exposure can shape subsequent immune responses through immune imprinting, potentially biasing antibody responses toward the previously encountered serotype and influencing the breadth of immunity generated against heterologous serotypes (*32*). This is particularly important for tetravalent dengue vaccines, which must generate balanced immunity against all four DENV serotypes despite heterogeneous pre-existing immune histories in the target population (*33, 34*). To determine whether serotype-specific pre-existing immunity alters responses to the tetravalent mRNA vaccine, four groups of mice were primed with monovalent DENV1, DENV2, DENV3, or DENV4 prM-E (Mono_D1prM-E_, Mono_D2prM-E_, Mono_D3prM-E_, or Mono_D4prM-E_) mRNA-LNP to establish distinct serotype-specific immune backgrounds. All groups subsequently received two Tet_prM-E_ mRNA-LNP boosts (**Fig. 7A**). A control group received Tet_prM-E_ mRNA-LNP for all three immunizations.

**Figure 7.**
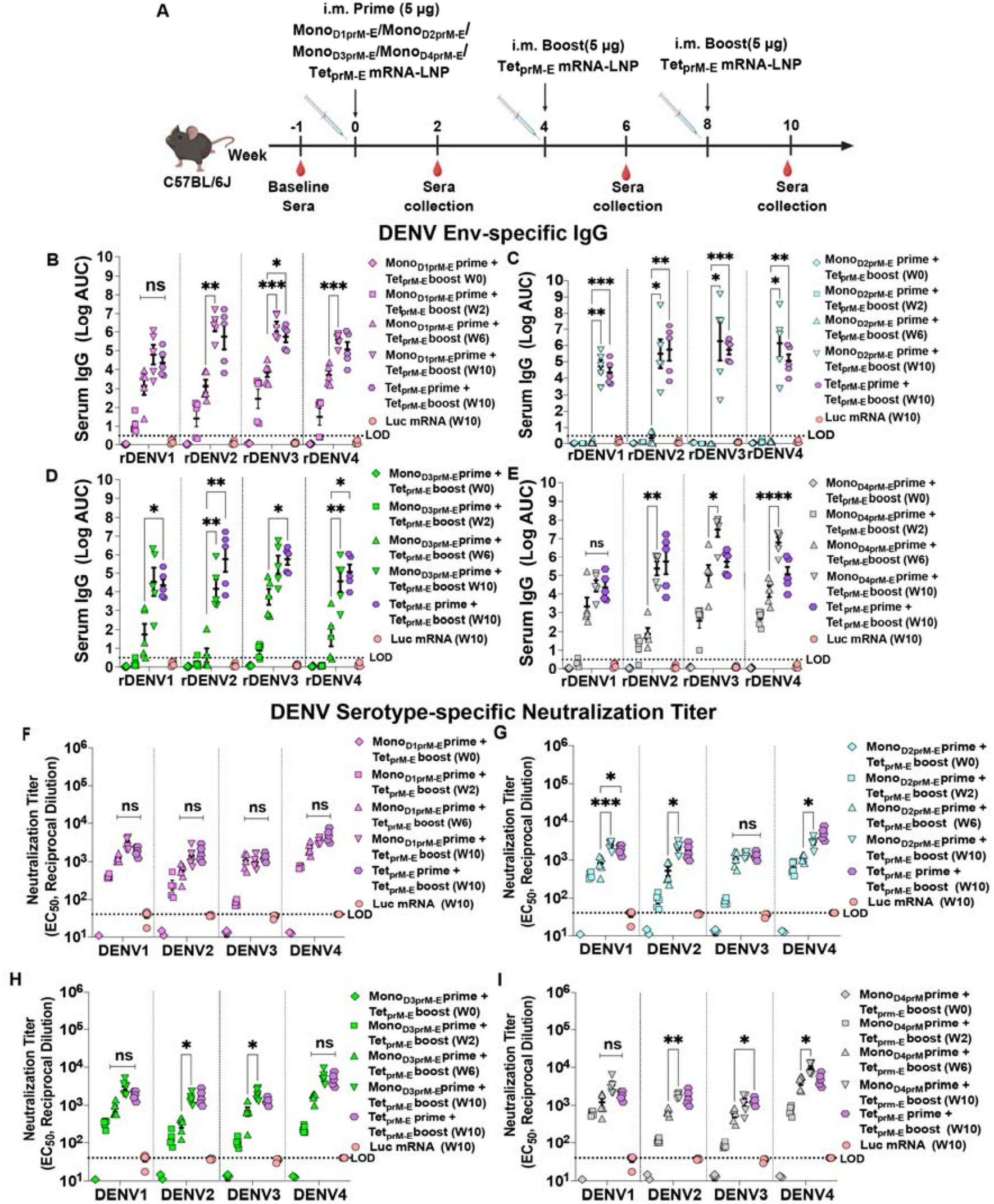
Evaluation of DENV serotype-specific immune imprinting following tetravalent vaccination. **(A)** Schematic of the DENV imprinting study. Five groups of C57BL/6J mice were separately primed intramuscularly (i.m.) at with 5 μg of DENV1, DENV2, DENV3, or DENV4 prM-E (Mono_D1prM-E_, Mono_D2prM-E_, Mono_D3prM-E_, or Mono_D4prM-E_) mRNA-LNP, or with tetravalent DENV prM-E (Tet_prM-E_) mRNA-LNP. All groups subsequently received 5 μg Tet_prM-E_ mRNA-LNP boosts at weeks 4 and 8. Serum was collected at baseline and at weeks 2, 6, and 10. Serum IgG binding responses to recombinant DENV1-4 envelope (env) antigens in mice primed with Mono_D1prM-E_ **(B)**, Mono_D2prM-E_ **(C)**, Mono_D3prM-E_ **(D)**, or Mono_D4prM-E_ **(E)** mRNA-LNP and boosted with Tet_prM-E_ mRNA-LNP. Responses are shown as log area under the curve (AUC) at weeks 0, 2, 6, and 10. Neutralizing antibody responses against DENV1-4 in mice primed with Mono_D1prM-E_ **(F)**, Mono_D2prM-E_ **(G)**, Mono_D3prM-E_ **(H)**, or Mono_D4prM-E_ **(I)** mRNA-LNP and boosted with Tet_prM-E_ mRNA-LNP. Neutralization titers are reported as EC_50_ reciprocal serum dilution. Tet_prM-E_ mRNA-LNP-primed mice and luciferase (luc) mRNA-LNP controls are shown for comparison. Dashed horizontal lines indicate the limit of detection (LOD), corresponding to a 1:40 serum dilution. Data are presented as mean ± SEM; n = 5 mice per group. Statistical significance was assessed using a mixed-effects model (REML) with the Geisser-Greenhouse correction, followed by Tukey’s multiple-comparisons test. ns, not significant; \**P* < 0.05, \*\**P* < 0.01, \*\*\**P* < 0.001, \*\*\*\**P* < 0.0001.

Serotype-specific env-binding IgG was measured longitudinally to determine whether monovalent priming restricted the breadth of antibody responses elicited by subsequent tetravalent vaccination. As expected, antibody binding was negligible at baseline, whereas monovalent priming preferentially increased responses to the corresponding serotype by week 2, confirming establishment of distinct serotype-specific immune backgrounds **(Fig. 7B-E)**. Following the first Tet_prM-E_ mRNA-LNP boost, responses broadened substantially by week 6 in most monovalent-primed groups, extending recognition to env from serotypes not represented during the initial monovalent immunization. Notably, this broadening was not uniform across priming backgrounds. Mono_D2prM-E_-primed animals showed limited expansion of env-binding at week 6 (Fig 7C), indicating that DENV2-specific pre-existing immunity was associated with delayed development of the tetravalent binding response.

Following the second Tet_prM-E_ mRNA-LNP boost, binding responses were maintained or further increased across all four serotypes in every priming group by week 10. In Mono_D2prM-E_-primed animals, responses increased significantly from week 6 to week 10 against DENV1-4 env antigens (**Fig. 7C**). Despite different priming histories, week 10 responses were generally comparable to those in mice receiving Tet_prM-E_ mRNA-LNP for all three immunizations, with no significant differences for most serotype-specific comparisons (**Fig. 7B-E**). Thus, serotype-specific priming altered the kinetics but did not persistently restrict the breadth or magnitude of E-binding responses following tetravalent boosting.

We next examined whether the antibody responses elicited following sequential tetravalent boosting were accompanied by broad functional neutralizing activity against all four DENV serotypes. Following monovalent priming, neutralizing activity was predominantly directed toward the corresponding priming serotype, consistent with the establishment of serotype-specific immunity. Subsequent Tet_prM-E_ mRNA-LNP vaccination broadened functional neutralization, with neutralizing activity against all four DENV serotypes detectable by week 6 and maintained or further increased at week 10 across the monovalent-primed groups **(Fig. 7F-I)**. Notably, in Mono_D2prM-E_-primed animals, the increase in total env-binding IgG was limited at week 6 **(Fig. 7C)**, yet functional neutralizing activity against all four DENV serotypes was already detectable at this time point, EC_50_ titers of 800 ± 170, 600 ± 150, 1100 ± 200, and 1100 ± 100 against DENV1, DENV2, DENV3, and DENV4, respectively **(Fig. 7G)**. This finding indicates that functional neutralization had broadened after a single tetravalent boost despite the comparatively modest increase in env-binding antibody responses. Following the second Tet_prM-E_ mRNA-LNP boost, robust neutralizing responses against all four serotypes were observed regardless of the initial monovalent prime. Importantly, by week 10, neutralization titers in monovalent-primed animals were generally comparable to those achieved following three sequential Tet_prM-E_ mRNA-LNP immunizations, with no significant differences for the majority of serotype-specific comparisons **(Fig. 7F-I)**. Together, these findings show that serotype-specific immune imprinting can shape the kinetics, but not the eventual breadth, of Tet_prM-E_ mRNA-LNP-induced immunity, highlighting its potential to overcome immune-history-dependent constraints and establish broad and balanced protection against the four dengue serotypes.

## Discussion

Our results establish Tet_prM-E_ mRNA-LNP vaccination as a strategy of generating balanced and functionally integrated immunity against dengue. Tetravalent vaccination induced neutralizing antibodies against DENV1-4, established CD4-dependent germinal center responses and durable B cell memory, and protective humoral immunity. Importantly, although prior monovalent immunity altered the response kinetics, repeated tetravalent vaccination ultimately generated four-serotype neutralization regardless of the priming serotype, supporting the capacity of this platform to broaden immunity across distinct pre-existing immune states.

Achieving effectively balanced immunity against all four DENV serotypes remains a major challenge in dengue vaccine development. Natural DENV infection typically induces long-lasting protection against the infecting serotype, whereas cross-protection against heterologous serotypes is transient (*35*). Importantly, waning cross-reactive antibodies may contribute to ADE during subsequent heterologous infection (*36*). Tetravalent live-attenuated vaccines have likewise shown serotype-dependent differences in immune responses and protection (*37–39*), with responses often dominated by cross-reactive rather than type-specific antibodies (*19*). Although higher neutralizing titers correlate with reduced dengue risk (*40, 41*), neutralization magnitude alone may not define protection; antibody specificity and functional quality are also important (*42*). Thus, evaluating tetravalent dengue vaccines requires consideration of not only the breadth and magnitude of responses across serotypes, but also their functional quality, specificity, and durability. Recently, efforts have been made to mutate the fusion loop and prM epitopes in replication-competent dengue viruses to reduce the elicitation of potential problematic cross-reactive antibodies (*43, 44*).

Tet_prM-E_ mRNA-LNP vaccine induced neutralizing activity against all four serotypes, although titers differed quantitatively. DENV4 showed the highest neutralization against mature virus, whereas DENV3 elicited the strongest env-binding IgG response, indicating that binding magnitude did not predict neutralizing potency. This is consistent with studies showing that potent human DENV-neutralizing antibodies frequently recognize conformational and quaternary env epitopes on intact virions, including broadly neutralizing env epitopes and serotype-specific epitopes that differ among DENV serotypes (*45–47*). Neutralization also varied with virion maturation, likely reflecting altered epitope exposure caused by incomplete prM cleavage (*48*). Consistent with our observations, a recent longitudinal study of 1,206 children showed that neutralizing titers measured against standard and mature DENV differed substantially in their relationship with protection; notably, 43% of dengue cases had standard-virus PRNT titers >100 compared with only 2% when mature virus was used, suggesting that mature-virus assays may more selectively capture functionally relevant neutralizing activity (*49*). These findings emphasize the importance of epitope specificity and accessibility, in addition to antibody magnitude, in shaping functional neutralization.

The strong GC response induced by Tet_prM-E_ mRNA-LNP vaccine provided a mechanistic basis for antibody maturation. GCs drive somatic hypermutation, clonal selection, and affinity maturation (*26, 50*). CD4+ T cell depletion during boosting markedly reduced Tfh cells and both total and env-specific GC B cell populations, demonstrating strong dependence on CD4+ T cell help. This pathway may be especially relevant in dengue, where antibodies that bind below the stoichiometric threshold required for neutralization can instead enhance infection through Fcγ receptor-bearing cells (*29, 51*). Thus, Tfh-dependent GC activity may contribute to the development of functionally mature antibody responses.

Most importantly, passive transfer provided direct evidence that the Tet_prM-E_ mRNA-LNP-induced humoral response was protective. Transfer of immune serum completely protected highly susceptible AG129 mice from lethal DENV2 challenge, prevented progressive weight loss, and substantially reduced acute viremia. Previous DENV mRNA vaccine studies demonstrated protection after monovalent vaccination (*7, 15*), while more recent work showed protective efficacy following tetravalent prM-E mRNA-LNP vaccination (*52*). Our findings extend these observations by showing that serum alone was sufficient to transfer protection. Although transient viremia remained detectable, sterilizing immunity was not required for protection in this model. Evaluation against additional serotypes remains limited by dengue mouse models, as many clinical isolates require highly virulent strains, high challenge doses, or mouse adaptation to produce reproducible disease (*53–55*).

The ADE analysis further defined the functional profile of vaccine-induced antibodies. No enhancement was detected across a broad concentration range, with ADE emerging only after dilution into the sub-neutralizing range. This behavior is consistent with the concentration-dependent nature of DENV ADE and published K562 data (*56*). Importantly, the emergence of ADE following extensive dilutions is not unique to vaccine-induced serum or indicative by itself of a poorly functional antibody response. Human sera with high DENV-neutralizing titers can similarly become enhancing when diluted to sub-neutralizing concentrations (*57*). Thus, the key finding is not the eventual appearance of ADE after extensive dilution, but the broad non-enhancing range extending through 1:10,000, together with complete protection after passive transfer.

Tet_prM-E_ mRNA-LNP also elicited env-specific cellular immunity, including IFN-γ-producing and polyfunctional IFN-γ+TNF-α+ CD8+ T cells with predominantly memory phenotypes. Multifunctional memory CD8+ T cell responses have also been observed after protective tetravalent dengue vaccination in humans (*58*). Moreover, mRNA-LNP vaccination can promote MHC-I presentation and antigen-specific CD8+ T cell priming with memory formation (*59, 60*). Unlike natural DENV infection and live-attenuated vaccination, where CD8+ responses preferentially target NS3, NS4B, and NS5 (*61, 62*), Tet_prM-E_ mRNA-LNP generated env-specific responses. Whether these cells contribute directly to protection remains to be determined.

Humoral immunity persisted beyond peak immunogenicity, with env-binding antibodies to all four serotypes detectable for six months and accompanied by a sizeable env-specific MBC compartment. These cells expressed the memory-associated markers CD73, CD80, and PD-L2 (CD273), supporting establishment of a phenotypically differentiated memory population (*63, 64*). Their persistence, together with the earlier CD4/Tfh-dependent GC response, supports establishment of a durable and phenotypically differentiated B-cell memory compartment. Systems-serology profiling further showed that this persistent response included multiple antibody isotypes and subclasses together with broad Fc-receptor-binding profiles. Such features may be relevant in dengue, where Fc characteristics and effector functions have been associated with protection from symptomatic disease (*65*). Although our data do not assign protection to specific Fc functions, they demonstrate that Tet_prM-E_ mRNA-LNP establishes humoral immunity extending beyond neutralization alone.

Finally, Tet_prM-E_ mRNA-LNP broadened immunity across distinct pre-existing serotype-specific immune histories. Sequential DENV exposures can preferentially recall memory generated by the first serotype and potentially bias subsequent responses (*18, 34*). In our study, prior monovalent priming altered the kinetics and magnitude of tetravalent responses but did not determine their final breadth. After completion of the tetravalent regimen, mice primed with each individual serotype developed neutralizing activity against DENV1-4. The DENV2-primed group was particularly informative: env-binding IgG expanded slowly, yet four-serotype neutralization was already detectable after the first tetravalent boost. Although the underlying clonal mechanisms remain unresolved, these findings indicate that pre-existing serotype-specific immunity biases the trajectory rather than irreversibly constraining the breadth of the Tet_prM-E_ mRNA-LNP response. This capacity to achieve four-serotype neutralization across monotypic immune backgrounds supports further evaluation of tetravalent mRNA-LNP vaccination in increasingly complex DENV exposure histories.

### Limitations and Future Direction

The findings of this study also identify several opportunities to further define and advance the Tet_prM-E_ mRNA-LNP vaccine. First, this tetravalent vaccine contains prM, an antigen that can elicit poorly neutralizing antibodies capable of mediating ADE. Although enhancement was not detected across a broad range of immune-serum concentrations, determining the contribution of prM-directed antibodies to the overall response will be important for further antigen optimization. We attempted to address this experimentally by generating an M-E construct lacking the pr domain; however, this construct did not produce detectable antigen expression (data not shown), indicating that additional engineering will be required to reduce prM antigenicity while preserving efficient prM-E processing and env expression. Second, the functional breadth demonstrated by neutralization was necessarily greater than that examined in the in vivo protection and ADE experiments, which were restricted by the availability of suitable serotype-specific experimental systems. Protection was established against lethal DENV2 challenge, whereas ADE was evaluated using DENV4; extending these analyses across serotypes will require appropriately validated challenge and enhancement models, a persistent constraint in dengue preclinical research. Finally, these studies were performed in murine models and provide a controlled framework for defining immunogenicity, durability, and the influence of pre-existing monotypic immunity, but the translational significance of these findings will require validation in non-human primates and ultimately humans. In parallel, epitope-resolved mapping of the neutralizing antibody response will be important for determining whether tetravalent breadth arises from serotype-specific antibodies, cross-reactive neutralizing populations, or a combination of both.

## Materials and methods

All dengue virus mouse infections were performed in an animal biosafety level 2 (ABSL2) facility with approval from the Yale Institutional Animal Care and Use Committee and Yale Environmental Health and Safety.

### Cells and viruses

Vero-81 and HEK 293T cells were cultured at 37° C and 5% CO_2_ in Dulbecco’s Modified Eagle medium (DMEM, Gibco, Cat#11-965-092) supplemented with 10% heat-inactivated fetal bovine serum (HI-FBS, Millipore-Sigma, Cat#F4135-500ML), and 1% 10,000 U/mL of penicillin–streptomycin (pen-strep) (Gibco, Cat#15140-122). C6/36 mosquito cells (ATCC, CRL-1600) were cultured at 32° C and 5% CO_2_ in Minimum Essential Medium (MEM, Gibco, Cat#11095-080) supplemented with 5% HI-FBS, (Millipore-Sigma, Cat#F4135-500ML), 1X MEM Non-Essential Amino Acids Solution (MEM NEAA, Gibco, Cat#11140-050), and 1% pen-strep (Gibco Cat#15140-122). K562 cells (ATCC, CRL-3343) maintained in suspension culture in RPMI-1640 supplemented with 10% FBS, 2 mM L-glutamine, and 1% pen-strep. Vero-furin cells were cultures at 37° C and 5% CO_2_ in DMEM (Gibco, Cat#1-965-092) supplemented with 10% HI-FBS, (Millipore-Sigma, Cat#F4135-500ML), and 1% pen-strep (Gibco, Cat#15140-122), and 2.5 µg/mL puromycin (Invivogen, Cat#ant-pr-1).

DENV1 isolate (DENV1/Vietnam/2008/Yale-DH144, Gene bank# OR389016.1), DENV2 isolate (DENV2/Indonesia/1984/Yale0-DH069, Gene bank#PX297372.1), DENV3 isolate (DENV3/Cambodia/2011/Yale-DH166, Gene bank#OR389033.1), and DENV4 isolate (DENV4/Venezuela/1990/Yale-DH127, Gene bank#OR389355.1) a gift from Nathan. Grubaugh from Yale School of Public Health. DENV-2 mouse-adapted strain S221 was a gift from Dr. Sujan Shresta from the La Jolla Institute for Immunology (*66*). DENVs were propagated in confluent monolayers of C6/36 cells grown in T175 flasks to get the mature and partially immature viruses. The culture medium was removed, and the cells were washed once with 1× PBS before inoculation with 2 mL of viral stock and 1 mL of Opti-MEM reduced-serum medium supplemented with GlutaMAX (Gibco, Cat#51-985-034). Cells were incubated with the viral inoculum for 1 hour at 32°C with 5% CO_2_. Following incubation, an additional 22 mL of Opti-MEM supplemented with 2% HI-FBS was added, and the infected cells were incubated for 4 days for DENV1-4 and 3 days for DENV2 mouse-adapted S221. Culture supernatants were collected, centrifuged at 500 × g for 5 min to remove cellular debris, aliquoted and stored at −80°C until use. For propagation of mature DENV1-4 viruses, Vero-furin cells were used because they overexpress the human protease furin, facilitating efficient prM cleavage and production of mature virions with reduced uncleaved prM (*23*). Briefly, the culture medium was removed from confluent Vero-furin monolayer grown in T175 flasks, and the cells were washed once with 1× PBS before inoculation with 1 mL of viral stock diluted in 2 mL of DMEM without FBS. Cells were incubated for 1 hour at 37 °C with 5% CO_2_, followed by additional 22 mL of DMEM supplemented with 2% HI-FBS, and incubated for 8 days for all four DENV serotypes. Culture supernatants were harvested, centrifuged at 500 × *g* for 5 min, aliquoted and stored at −80 °C until further use. Virus titers were determined by focus-forming assay (FFA), as described previously (*67*). Experiments with DENV serotypes were conducted under biosafety level 2 (BSL2) containment at Yale School of Medicine with Institutional Biosafety Committee approval.

### Vaccine design

mRNAs were generated as described (*68*). Plasmids synthesized by GenScript were linearized and used as templates for transcription with T7 RNA polymerase (MEGAscript, Ambion). The plasmids encoded firefly luciferase (luc) and codon-optimized DENV prM-E sequences, including the corresponding signal peptide, for DENV1 (NC_001477 DENV1), DENV2 (MW946475_Tonga’74), DENV3 (NC_001475), and DENV4 (NC_002640). All mRNAs contained a 101-nucleotide poly(A) tail. For nucleoside modification, uridine-5′-triphosphate was replaced with N1-methylpseudouridine-5′-triphosphate (m1Ψ; TriLink) during transcription. Capping of mRNA was performed in concert with transcription through the addition of a trinucleotide cap1 analog, CleanCap (TriLink). Cellulose-based purification of the mRNAs was performed as described (*68*). The integrity of mRNAs was then assessed on an agarose gel before storing at −20°C.

Cellulose-purified m1Ψ-modified mRNAs encoding DENV prM-E or luc were formulated into lipid nanoparticles (LNPs) using microfluidic mixing of an organic phase containing lipids and an aqueous phase containing mRNA. The lipids were dissolved in ethanol and mixed at a molar ratio of 50:38.5:10:1.5 (SM102: Cholesterol: DSPC:DMG-PEG), and mRNA was diluted in 50 mM citrate buffer of pH 4 at a concentration of 129 ug/mL. Both phases were mixed at a volumetric ratio of 1:3 (ethanol: citrate buffer) using a microfluidic mixing device (NanoAssembler Ignite, Cytiva). Lipids and mRNA were mixed at a weight ratio of 40:1 for mRNA-LNP formulations. The LNPs were then dialyzed against 1x PBS buffer in 12-14 kDa dialysis membrane for 2 hours. The LNP formulations were characterized for their size, polydispersity index, and zeta potential using Zetasizer Pro ZS (Malvern Panalytical). The encapsulation efficiency of mRNA in LNPs was analyzed using the Quant-iT RiboGreen assay. The mRNA-LNPs were stored at −80°C at an mRNA concentration of approximately 0.5 μg/μL.

### Animal studies

The vaccination and viral challenge protocols were approved by the Institutional Animal Care and Use Committee (IACUC) at Yale School of Medicine (Animal protocol number: 20520).

### Vaccination for immunogenicity assessment

The tetravalent DENV prM-E (Tet_prM-E_) mRNA-LNP formulation was prepared by combining equal amounts of DENV1-4 monovalent prM-E mRNA-LNPs (1.25 µg of each; total dose of 5 µg). C57BL/6J mice used in this study were bred at Yale School of Medicine. Eight-week-old male and female C57BL/6J mice were primed intramuscularly (i.m.) with a 5 µg Tet_prM-E_ mRNA-LNP and received addition two i.m. 5 µg booster immunizations at 4-week intervals. Control mice received 5 µg of luc mRNA-LNP on same schedule. Baseline sera were collected before priming, and sere were collected 2 weeks after each immunization for immunogenicity assessment.

### CD4^+^ T cell depletion and germinal center responses

C57BL/6J mice were primed with a 5-µg dose of Tet_prM-E_ mRNA-LNP on day 0. For *in vivo* depletion of CD4+ T cells, mice received 125 µg of anti-mouse CD4 monoclonal antibody (clone GK1.5, bioXcell, Cat**#**BE0003-1) or isotype-matched control antibody (rat IgG2b, clone LTF-2, bioXcell, Cat**#**BE0090) by intraperitoneal (i.p.) injection in 50 µL sterile 1X PBS on day 6. Mice received 5-µg Tet_prM-E_ mRNA-LNP i.m. boost on day 7 and a second i.p. dose of depleting or control antibody on day 8. On day 14, mice were humanely euthanized and vaccine-draining inguinal lymph node (dLN) and spleen were collected. Depletion efficacy was assessed by flow cytometry using non-competing anti-CD4 antibody (clone RM4-4, BioLegend, Cat#11600).

### Sequential vaccination and immune imprinting

To establish DENV serotype-specific immune imprinting, mice were divided into four groups and primed i.m. with a 5 µg dose of Mono_D1prM-E_, Mono_D2prM-E_, Mono_D3prM-E_, or Mono_D4prM-E_. All groups subsequently received two i.m. 5-µg of Tet_prM-E_ mRNA-LNP boosts. An additional control group received Tet_prM-E_ mRNA-LNP for all three immunizations at the same dose. Baseline sere were collected before vaccination, and 2 weeks after each immunization for immunogenicity assessment.

### Antibody durability and memory B cell responses

Mice were immunized i.m. with Tet_prM-E_ mRNA-LNP using a three-dose regimen of one prime and two boosts, 5 µg per immunization. To evaluate the durability, terminal collections were performed at weeks 10, 14, 22, and 34 after the initial immunization. At each time point, designated mice were euthanized, and serum and spleens were collected. Serum was used to assess DENV-specific antibody responses, and splenocytes were used to characterize DENV-specific MBC responses.

### Dengue virus 2 challenge study

Six-week-old male and female AG129 (IFN-α/β/γR^−/−^) mice, originally provided by Dr. Sujan Shresta and bred in house, were used to evaluate the protective efficacy of sera from Tet_prM-E_ mRNA-LNP-vaccinated mice against lethal DENV2 challenge. Mice were passively immunized 18 hours before viral challenge with either sera from Tet_prM-E_ mRNA-LNP-vaccinated mice (350 µL/mouse), PBS (350 µL/mouse), or EDE1-C10 antibody (100 µg/mouse in 350 µL of PBS) administered intraperitoneally. Mice were challenged retro-orbitally with 1 × 10^6^ FFU of mouse-adapted DENV2 strain S221. Body weight and clinical signs of disease were monitored daily. Mice were humanely euthanized upon reaching the predefined humane endpoint of ≤80% of their initial body weight or upon developing significant clinical signs of disease. Serum samples were collected daily for up to 10 days after challenge, and viral titers were determined using a focus-forming assay (FFA).

### In vitro transfection and viral protein detection

HEK293T cells were transfected with TransIT-mRNA (Mirus Bio) according to the manufacturer’s instructions. For all constructs except DENV2 prM-E, 0.21 µg mRNA was mixed with 0.23 µl TransIT-mRNA Reagent and 0.15 µl Boost Reagentin 17 µl of serum-free medium and added to 7 × 10⁴ cells seeded the previous day in 183 µl of complete medium. For DENV2 prM-E mRNA, 0.23 µg mRNA was mixed with 0.7 µl of each TransIT-mRNA and Boost Reagent in 18 µl of serum-free medium; 17 µl was added to cells under the same conditions. At 15-24 hours after transfection, cells were lysed on ice for 1 hour in RIPA buffer (Thermo Scientific). Each construct was transfected in triplicate to increase lysate yield.

Whole-cell lysates were analyzed for DENV env and prM protein expression by non-denaturing SDS-PAGE western blot. Samples were heated at 99 °C for 5 min, mixed with 4× Laemmli buffer (Bio-Rad), and separated on 4-15% polyacrylamide Mini-PROTEAN TGX precast gel (Bio-Rad) for 20 min at 80 V, followed by 60 min at 120 V. Proteins were transferred to a nitrocellulose membrane using a Trans-Blot Turbo Transfer System and 0.2-µm Nitrocellulose Transfer Packs (Bio-Rad).

For DENV env protein detection, membranes were blocked overnight at 4 °C in 5% BSA in PBS containing 0.5% Tween-20 (PBS-T), then probed for 2 hours at 37 °C with week-10 sera from Tet_prM-E_ mRNA-LNP vaccinated mice (1:5,000). Membranes were incubated with HRP-conjugated goat anti-mouse IgG secondary antibody (SeraCare; 1:5,000) for 1 hour at room temperature (RT). Both antibodies were diluted in 3% BSA in PBS-T. Signal was developed using Clarity Western ECL substrate (Bio-Rad) and imaged using the ChemiDoc imaging system (Bio-Rad). Washing steps were performed with PBS-T. DENV prM detection was performed identically, except that membranes were blocked for 1 hour at RT and incubated with primary antibody overnight at 4 °C.

### ELISA

DENV E protein-specific serum IgG responses were measured by enzyme-linked immunosorbent assay (ELISA). Half-area 96-well plates (Corning, Cat#3690) were coated overnight at 4°C with recombinant DENV1-4 env protein dimers at 1 μg/mL in 0.1 M carbonate buffer (pH 9.6; 25 μL/well), DENV1 env protein (strain US/Hawaii/1944; His-tagged dimer; Sino Biological, Cat#40531-V08H1-10) and in-house DENV2-4 env protein dimers were used as capture antigens. Plates were washed once with tris buffer saline containing 0.2% Tween-20 (TBS-T) and blocked with 5% non-fat milk in TBS-T (50 μL/well) for 2 hours at RT with gentle rocking. Serially diluted sera prepared in blocking buffer were added at 25 μL/well and incubated overnight at 4°C. Dilutions began at 1:3,000, and each sample was tested against all four DENV serotypes. Plates were washed three times with TBS-T and incubated with horseradish peroxidase (HRP)-conjugated goat anti-mouse IgG (SeraCare, Cat#5220-0341; 1:1,500; 25 μL/well) for 2 hours at RT. After four washes, 25 μL of TMB substrate (SeraCare, Cat#5120-0083) was added, and reaction was developed in dark at RT, and stopped with 25 μL of 0.16 M sulfuric acid. Absorbance was measured at 450 nm. The ELISA limit of detection (LOD) was defined as the mean blank signal + 3 × SD of the blank.

### Focus-reduction Neutralization Test (FRNT)

Neutralization assays were performed as previously described (*67*). Vero-81 cells were seeded at 200,00 cells/well in 96-well plates one day before the focus reduction neutralization test (FRNT). Sere were heat-inactivated at 56°C for 30 min, diluted 1:20 in infection medium (DMEM with 2% HI-FBS and 1% pen-strep), and serially diluted 1:3 for eight dilution points. DENV stocks were diluted to 3,500 FFU/mL for DENV1-DH144 and DENV2-DH069, and 4,000 FFU/mL for DENV3-DH166 and DENV4-DH127. Equal volumes (30 μL) of virus and diluted serum were mixed and incubated at 37°C for 1 hour. Medium was removed from the Vero-81 cells, 50 μL of each serum-virus mixture was added, and plates were incubated at 37°C for 1 hour. Overlay medium (125 μL/well; Opti-MEM with 2% HI-FBS, 1% pen-strep, and 1% methylcellulose) was then added. Plates were incubated at 37°C for 52 hours for DENV4 or 57 hours for DENV1-3. Cells were fixed with 10% formaldehyde, permeabilized with 1X permeabilization buffer (Invitrogen, Cat#00-8333-56;), and blocked for 10 min in same buffer containing 5% nonfat dry milk. Cells were incubated with pan-flavivirus murine monoclonal antibodies 4G2 and 2H2 (1:500), followed by HRP-conjugated anti-mouse IgG (H+L) (SeraCare, Cat#5220-0341; 1:1,500). Foci were developed using TrueBlue peroxidase substrate (SeraCare, Cat#5510-0030), air-dried, imaged on a CTL ImmunoSpot reader, and quantified using Fiji-image J software.

### T cell stimulation

DENV2 CD4 MegaPool (MP) and DENV2 CD8 MP are synthetic peptide pools based on experimentally validated DENV epitopes (*69–71*). DENV2-specific MPs consisted of 15-mer peptides overlapping by 10 amino acids and spanning the prM, env, and NS1 proteins from the DENV2 PL046 strain. Peptides were resuspended in DMSO and combined into protein-specific pools containing 33 prM, 99 env, and 70 NS1 peptides, then lyophilized to minimize the final DMSO concentration during stimulation.

For T-cell stimulation assays, spleens were collected at week 10 (week 10) from Tet_prM-E_ mRNA-LNP or luc mRNA-LNP vaccinated mice following the three-dose regimen. Spleens were processed into single cell suspensions and cryopreserved until use. Splenocytes were thawed in RPMI 1640 medium (Corning) supplemented with 10% FBS, 1% Pen-strep, 42 mM HEPES (Sigma-Aldrich), and benzonase (0.1 KU/μL; 5 μL/10 mL of medium). Splenocytes were plated at 1-2 × 10^6^ cells/well in 96-well U-bottom plates and stimulated with DENV2-specific MPs (1 mg/mL). DMSO served as negative control, and phorbol 12-myristate 13-acetate (PMA; 100 μg/mL) plus ionomycin (1 mg/mL) (PMI) served as the positive control. After 1 hour, GolgiPlug (BD Biosciences, Cat#555029) was added for an additional 4 hours.

Cells were washed and stained with LIVE/DEAD Blue viability dye (Invitrogen) for 20 min, followed by anti-mouse Fc-receptor blocking antibody (BioLegend) for 10 min. Surface staining was performed for 20 min at 4°C using CDε-BV480 (Thermo Fisher Sci. Cat#414-0031-82), CD4-BUV395 (BD Bioscience, Cat#563790), CD8-BV510 (BioLegend, Cat#100752), CD44-BV785 (BioLegend, Cat#103059), CD62L (L-selectin)-APC-eFlour 780 (Thermo Fisher Sci. Cat#47-0621-82). Cells were then fixed with 4% paraformaldehyde for 10 min and permeabilized with 0.1% saponin in PBS and stained intracellularly for 30 min at RT with IL-2-BV711 (BioLegend, Cat#503837), TNF-α-APC (BioLegend, Cat#506308), IFN-γ-FITC (Cytek Biosciences, Cat#72216), Granzyme B-PE-Dazzle 594 (BioLegend, Cat#101302).

Samples were acquired on a Cytek Aurora spectral flow cytometer (Cytek Biosciences) and analyzed using FlowJo, OMIQ (Dotmatics), and GraphPad Prism. Antigen-specific responses were calculated after subtraction of the corresponding DMSO background. A response was considered positive when the background-subtracted value exceeded the limit of the machine (LOM) and the stimulation index was >2. The stimulation index was calculated as the response to the indicated DENV2 peptide pool divided by the DMSO control responses.

### Systems serology

Serum antibody responses were profiled using a multiplexed bead-based systems serology assay, following a previously described method (*72*). Briefly, carboxylated MagPlex microspheres (Diasorin) were covalently linked with DENV E-derived antigens and control proteins via NHS-ester linkages by incubating with EDC and sulfo-NHS (Thermo Fisher). Antigens and controls were sourced from Sino Biological: DENV-E (type 1, US/Hawaii/1944), Dimer (Cat#40531-V08H1), DENV-E Protein (Cat#40531-V08H), DENV-E (type 1, US/Hawaii/1944), Domain III (Cat#40531-V08B), DENV-E (type 2, New Guinea C/PUO-218 hybrid), Domain III (Cat#40471-V08Y1), DENV-E (type 4, Philippines/H241/1956), Dimer (Cat#40533-V08H1), DENV-E (type 3, Philippines/H87/1957), aa 579-678 (Cat#40532-V08B3), DENV-E (type 4, Philippines/H241/1956), aa 579-679 (Cat#40533-V08B), HCMV gB (Cat#10202-V08H1), HSA (Cat#10968-HNAY), and Ebola gP (Cat#40459-V08H). Linear ranges of detection for each antibody isotype, subclass, and FcγR-binding antibody were determined using randomly selected specimens prior to running the cohort. Dilutions used were 1:200 for Total IgG, IgG1, IgG2a, IgG2b, and IgG2c, 1:100 for IgG3, IgA, and IgM, and 1:400 for Fcγ2b, Fcγ3, and Fcγ4. All serum samples were diluted in 1X PBS pH = 7.4. The samples were incubated with the antigen-coated bead slurry in 1X Assay buffer (1X PBS pH = 7.4, 0.1% BSA, and 0.02% Tween-20) overnight at 4°C in non-binding 384-well plates (Greiner Bio One), with continuous shaking at 800 rpm. Following incubation, unbound antibodies were removed using a magnetic 384-well HydroSpeed Plate Washer (Tecan) with 1X Assay buffer. Antibody isotypes and subclasses were detected with diluted phycoerythrin (PE)-conjugated secondary antibodies against Total IgG1, IgG2a, IgG2b, IgG2c, IgG3, IgA, and IgM (Southern Biotech) in 1X Assay buffer for 1 hour at room temperature with continuous shaking. Complexes were then washed with 1X Assay buffer using the HydroSpeed Plate Washer. After the final wash, beads were resuspended in 30 μL of 1X xMAP Sheath Fluid PLUS Additive (Thermo Fisher) and protected from light until analysis (see below).

FcγR-binding activity was assessed by biotinylating AVI-tagged FcγR2b, FcγR3, and FcγR4 custom synthesized by the Duke Protein Production Facility using the BirA500 kit (Avidity LLC). Following biotinylation, the receptors were complexed with PE-streptavidin (Agilent Technologies) at a 1:4 molar ratio. The PE-streptavidin-FcγR were incubated with the serum-incubated bead slurry (diluted at 1:100 in 1X Assay buffer) for 1 hour at room temperature with continuous shaking. Complexes were washed with 1X Assay buffer. After the final wash, beads were resuspended in 30 μL 1X xMAP Sheath Fluid PLUS Additive (Thermo Fisher) and protected from light until use.

All binding measurements were captured same-day using the Luminex xMAP Intelliflex System (Thermo Fisher) with readouts of median fluorescence intensity (MFI) reported in arbitrary units (AU). All samples were run in duplicate, and the means of the technical replicates were reported. A region scatter/gating strategy is shown in **Fig. S7**. Antigen-binding responses were subsequently expressed as averaged MFI or MFI fold change over naïve samples, with the naïve response normalized to a value of 1.

### Multiplex Immunofluorescence imaging

Tissue sections were processed for multiplex imaging using the Cell DIVE Multiplex Imaging Solution (Leica Microsystems), as previously described (*73*). Slides were rehydrated three times for 5 min in PBS, then blocked and permeabilized for 1 hour in PBS containing 0.2% BSA, 0.05% sodium azide, 0.3% Triton X-100, and 10% FBS. Slides were placed into ClickWells (Leica Microsystems) and washed twice with PBS containing 0.01% Tween-20 (PBST), stained with 1 µg/ml DAPI for 15 min and washed. After addition of 1 mL 50% glycerol, initial autofluorescence images were acquired.

For biomarker staining, sections were incubated with antibodies (1:100 in 3% BSA) for 1 hour, washed twice with PBST for 2 min, stained with DAPI for 2 min, and imaged after addition of 1 mL glycerol. Following imaging, slides were washed and fluorophores were inactivated for 15 min with 100 mM NaHCO₃ and 3% H_2_O_2_ in ddH_2_O, followed by three PBST washes. Sequential rounds of biomarker staining, imaging, and dye inactivation were performed for up to three rounds until imaging was completed. Images were acquired using the Leica Cell DIVE Multiplex Imaging Solution and analyzed using HALO AI software (Indica Labs). Antibodies included Alexa Fluor 488 anti-mouse IgD (clone 11-26c.2a, BioLegend, Cat# 405718), Alexa Fluor 647 anti-mouse GL7 (clone GL7, BioLegend, Cat#144606), Alexa Fluor 488 anti-mouse B220 (clone RA3-6B2, Invitrogen, Cat#53-0452-82), PE anti-mouse CD4 (clone GK1.5, BioLegend, Cat#00408), Alexa Fluor 647 anti-mouse PD-1 (clone RMP1-30, BioLegend, Cat#109118), and PE mouse anti-BCL6 (clone K112-91, BD Biosciences, Cat#561522).

### Antibody-dependent enhancement (ADE) assay based on luminescence

ADE was assessed using K562 cells, as previously described (*44*). Cells were seeded in black, flat-bottom 96-well plates at 1 × 10^5^ cells/well in 50 µL of infection medium (RPMI-1640 with 2% FBS, 2 mM L-glutamine, and 1% pen-strep). Sera were serially diluted fourfold across eight dilutions begining1:80 dilution. DENV4-nLuc virus was diluted in the same medium to 5,000 FFU/well. Equal volumes of diluted serum and virus (25 µL each) were combined and incubated at 37°C for 1 hour. Cells plus virus without antibody served as the normalization control. Plates were incubated for 24 hours at 37°C with 5% CO_2_. Following incubation, Nano-Glo Luciferase Assay reagent (Promega) was added according to the manufacturer’s instructions and Luminescence was measured using a compatible luminometer. ADE was expressed as the fold change in luminescence relative to the no-antibody control.

### Flow cytometric analysis of T follicular helper cells, germinal center B cells, memory B cells

Mice were humanely euthanized on day 14, and the spleen and vaccine-draining inguinal lymph node (dLN) were collected. Single-cell suspensions were prepared by mechanical dissociation through 70-µm cell strainer, followed by ACK lysis buffer for red blood cells lysis for 1 min. Cells were washed, resuspended in FACS buffer, stained with Zombie NIR Fixable Viability Dye (BioLegend, Cat#423105, 1:100), and blocked with anti-mouse CD16/CD32 Fc Block (BioLegend, Cat#101302; 1:100) for 10 min at 4 °C prior to surface staining. Data acquired on a BD LSR Fortessa Flow Cytometer and analyzed using BD FACS Diva software (v.9.3.1) and FlowJo (v.10.10.0).

For Tfh-cell analysis, cells were stained with CD3-FITC (BioLegend, Cat#100203), CD4-Pacific Blue (BioLegend, Cat#116007), PD-1 (CD279)-PE/Cyanine7 (BioLegend, Cat#109109), and CXCR5 (CD185)-Brilliant Violet 711 (BioLegend, Cat#145529) for 30 min at 4 °C. Tfh cells were identified as PD-1+CXCR5+ cells within viable CD3+CD4+ T-cell population.

For GC B-cell analysis, DENV3 E-dimer protein was biotinylated using EZ-Link Sulfo-NHS-LC-Biotin (ThermoFisher, Cat#A39257) and conjugated with streptavidin-PE (Prozyme, Cat#PJRS25) and streptavidin-APC (Prozyme, Cat#PJ27S) at a 4:1 molar ratio at 4 °C to generate dual color DENV3 env-dimer probes. Cells were incubated with this probe for 30 min at 4°C, followed by staining with CD19-Spark PLUS UV395 (BioLegend, Cat#115587), B220-BUV737 (BD Biosciences, Cat#612839), CD3-Brilliant Violet 510 (BioLegend, Cat#100233), CD95 (Fas)-FITC (BioLegend, Cat#152605), GL7-Pacific Blue (BioLegend, Cat#144613), IgD-Brilliant Violet 711 (BioLegend, Cat#405731), and CD38-Alexa Fluor 700 (BioLegend, Cat#102742). GC B cells were defined as IgD^loCD38^loGL7^+^CD95^+^ cells, and DENV3 env-specific GC B cells as dual PE^+^APC^+^ probe-binding cells.

For MBCs, cells were stained with the same dual-color DENV3 env-dimer probes, followed by CD19-Spark PLUS UV395 (BioLegend, Cat#115588), CD3-FITC (BioLegend, Cat#100203), B220-BUV737 (BD Biosciences, Cat#612839), GL7-Pacific Blue (BioLegend, Cat#144614), CD38-Alexa Fluor 700 (BioLegend, Cat#102742), CD138-Brilliant Violet 510 (BioLegend, Cat#142521), CD80-PE/Cyanine7 (BioLegend, Cat#104733), PD-L2 (CD273)-PE/Dazzle 594 (BioLegend, Cat#107215), and CD73-Brilliant Violet 605 (BioLegend, Cat#127215). Memory-like B cells were defined as GL7⁻CD138⁻CD38^+^, and DENV3 env-specific cells as dual PE^+^APC^+^ probe-binding cells. Phenotypic characterization was based on CD273, CD80, and CD73 expression.

### Statistical Analysis

Statistical significance among groups was evaluated using two-way ANOVA followed by Tukey’s multiple comparisons test for ELISA and neutralization assays. For flow cytometry data, comparisons between two groups were performed using a two-tailed Mann-Whitney U test. Comparisons among multiple groups were performed using the Kruskal-Wallis test followed by Dunn’s multiple comparisons test. When two factors were analyzed with repeated measurements, statistical significance was assessed using a mixed-effects model (REML) with the Geisser-Greenhouse correction, followed by Tukey’s multiple-comparisons test. Data from the challenge study were analyzed using a mixed-effects model (REML) with the Geisser-Greenhouse correction, followed by Dunnett’s multiple comparisons test. Data from the ADE study were analyzed using Two-way Anova, followed by Dunnett’s multiple comparison test. All data are presented as mean ± SEM. *P* values < 0.05 were considered statistically significant (ns, not significant; \**P* < 0.05, \*\**P* < 0.01, \*\*\**P* < 0.001, and \*\*\*\**P* < 0.0001). *P* values were adjusted for multiple comparisons where applicable. All statistical analyses were performed using GraphPad Prism (version 11).

For the principal component analysis (PCA), all systems serology features were scaled across timepoints, and the PCA was generated using the R Studio packages dplyr, factoextra, ggfortify, and ggrepel. Negative response controls such as Ebolavirus glycoprotein, human cytomegalovirus glycoprotein b, and human serum albumin were excluded from the scaling. Composite profiles were scattered on principal component axes 1 and 2 (PC1 and PC2) and colored by week after vaccination. Ellipses were added to show the 95% confidence interval of the group scattering. A bitplot of the features contributing to the influence along PC1 and PC2. The top 8 features were shown, along with their vector influence.

## Acknowledgement/Funding

D.R.M is funded by a Hanna H. Gray Fellowship from the Howard Hughes Medical Institute.

DW is partially supported by P01 AI106695.

R.P.M. is funded by the U19AI135995.

## Author contributions

Conceptualization: D.R.M.

Methodology: F.A., Y.R.D., J.V., K.S.L, E.B., J.P., Z.Y., N.J., H.M., S. T., and S.S.

Investigation: F.A., Y.R.D., J.V., K.S.L, E.B., J.P., Z.Y., H.M., S.T., and S.S.

Visualization: F.A., Y.R.D., J.V., K.S.L, Z.Y., D.R.M.

Funding acquisition: D.R.M.

Project administration: D.R.M.

Supervision: D.R.M., N.P., L.V.T., D.W., and R.P.M.

Writing – original draft: F.A.

Writing – review & editing: F.A. Y.R.D., J.V., K.S.L, E.B., J.P., Z.Y., H.M., S.T., S.S., C.W., N.J., R.P.M., D.W., L.V.T., N.P., and D.R.M.

