## Supplemental materials for "Tetravalent dengue mRNA vaccination orchestrates balanced four-serotype immunity with durable memory and protection in mice"

**Supplemental results:**

**Dengue immunogen down-selection for a tetravalent mRNA vaccine**

We first focused on identifying immunogenic dengue virus antigen when delivered as mRNA-LNPs (lipid nanoparticles) before expanding our vaccination strategy as a tetravalent vaccine. As an initial model serotype, we focused on dengue virus serotype 2 (DENV2) structural proteins. We designed three constructs using MW946475_Tonga'74 DENV2 genome sequence: capsid (C), wild type precursor membrane (prM)- envelope (env): termed as non-stabilized prM-E, and a mutant prM-E containing the A259W and T262R substitution in env: named as stabilized prM-E, each including the corresponding signal peptide **(Fig. S1A)**. The A259W and T262R substitution were introduced to stabilization of env dimers (*21*). Mice were immunized with monovalent DENV2 mRNA-LNP constructs **(Fig. S1B)**. A negative control group was immunized with luciferase (luc) mRNA-LNP. Mice received one prime immunization followed by two booster immunizations at the indicated time points, and serum IgG responses were evaluated by ELISA against DENV2 whole virions. **Comparison of the area under the curve (AUC) values showed that the non-stabilized DENV2 prM-E construct was the most immunogenic and elicited the highest DENV2-specific IgG responses (Fig. S1C).** Thus, the non-stabilized prM-E construct was selected as the antigen for subsequent vaccine development and introduced as prM-E for rest of the study.

We next evaluated the immunogenicity of the individual serotype-specific prM-E constructs. Four groups of mice (n = 10 per group) were separately immunized with 5 µg of monovalent DENV1-4 prM-E (Mono_D1prM-E_, Mono_D2prM-E_, Mono_D3prM-E_, or Mono_D4prM-E_) mRNA-LNP. A negative control group (n=10) was immunized with luc mRNA-LNP. Mice were immunized using a prime-boost-boost regimen **(Fig S2A)**. Mouse vaccine serum samples were then evaluated for DENV serotype-specific antibody responses by whole-virion ELISA, with responses quantified as AUC values. Each monovalent prM-E formulation elicited antibody responses against its corresponding DENV serotype, as demonstrated by increased whole-virion ELISA AUC values compared with the luc mRNA-LNP control **(Fig S2B).** To determine whether antibodies elicited by monovalent prM-E vaccination were functionally neutralizing, we selected sera from Mono_D2prM-E_-immunized mice for further evaluation against the corresponding DENV2 serotype by focus reduction neutralization test (FRNT) (**Fig S2C**). We have used the standard virus stock growing in Vero-81 cells for this test. Mono_D2prM-E_ vaccination elicited robust DENV2-neutralizing antibody responses, with a mean EC₅₀ titer of **4,800 ± 510,** which is significantly higher compared to the luc mRNA-LNP control group **(*****P* < 0.0001**; **Fig. S2D**).

To determine whether these vaccine-elicited antibodies were sufficient to confer protection in vivo, sera from Mono_D2prM-E_ or luc mRNA-LNP-immunized donor mice were passively transferred into type I interferon receptor knock out (IFNAR1) mice prior to DENV2 lethal challenge with a DENV2 mouse-adapted strain S221. Mice received of Mono_D2prM-E_ mRNA-LNP-immunized sera exhibited 100% survival and were protected from disease-associated weight loss following DENV2 lethal challenge **(Fig. S2E, F)**. Together, these findings supported the selection of prM-E as the vaccine antigen and demonstrated the immunogenicity of the individual monovalent prM-E mRNA-LNP constructs prior to development of the tetravalent formulation.

**Supplementary figure legends:**
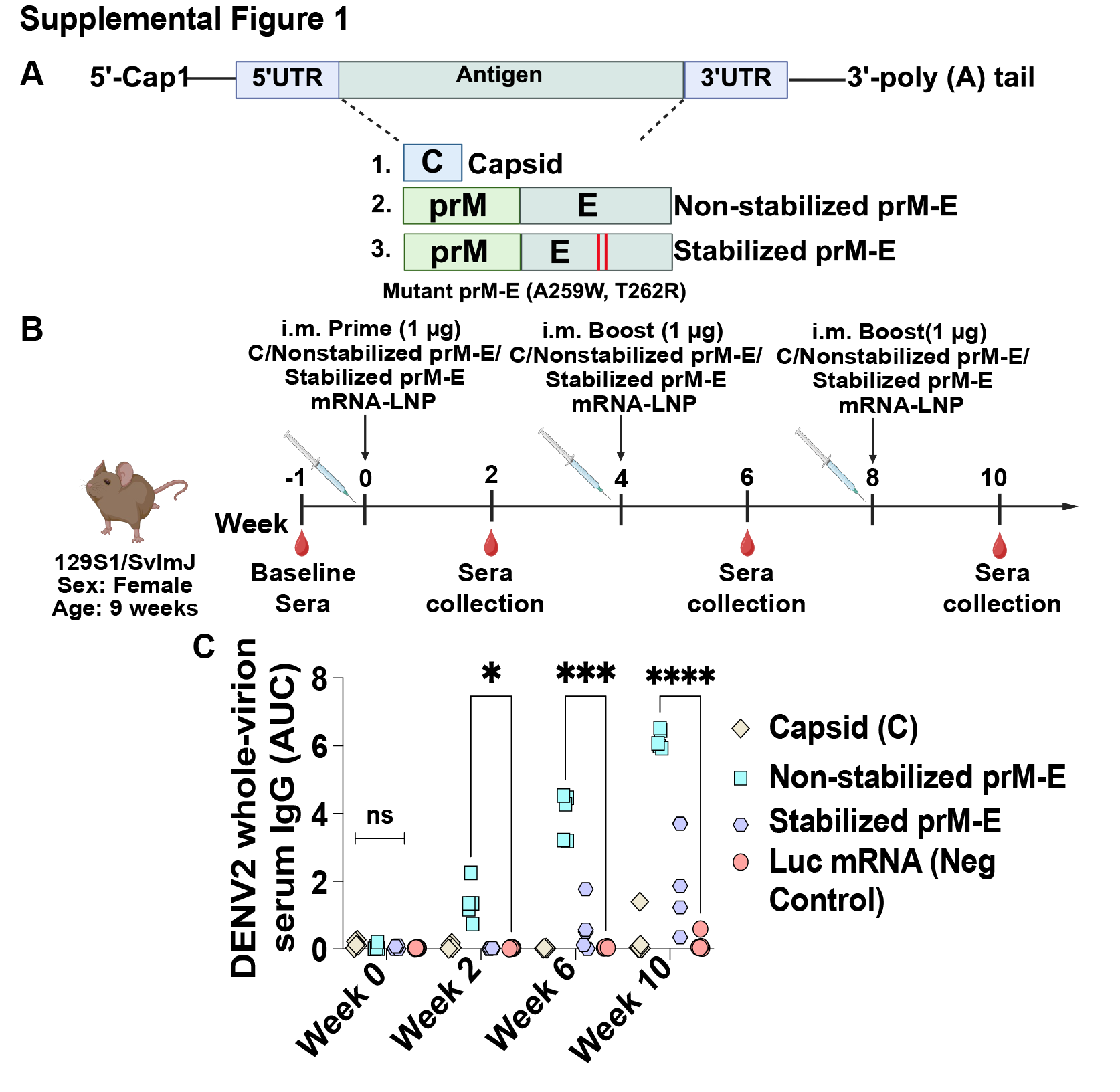

**Supplementary Figure 1:** **Down selection of DENV mRNA-LNP immunogens. (A)** Schematic of the mRNA constructs evaluated for antigen selection. Constructs encoded DENV2 capsid (C), non-stabilized prM-E, or stabilized prM-E containing the A259W and T262R substitutions. **(B)** Schematic of the immunization and serum collection schedule. Nine-week-old female 129S1/SvImJ mice (n = 5 per group) were immunized intramuscularly (i.m.) with 1 μg of monovalent DENV2 capsid or non-stabilized prM-E or stabilized prM-E mRNA-LNP at weeks 0, 4, and 8. Baseline serum was collected before prime, and post-immunization sera were collected at two weeks after each vaccination. **(C)** DENV2 whole-virion-specific serum IgG responses measured by ELISA at weeks 0, 2, 6, and 10 in mice immunized with capsid, non-stabilized prM-E, or stabilized prM-E mRNA-LNP. Luciferase (luc) mRNA-LNP-immunized mice served as negative controls. Antibody responses are expressed as area under the curve (AUC). Comparisons among multiple groups were performed using the Kruskal-Wallis test followed by Dunn’s multiple-comparisons test. ns, not significant; *P < 0.05, ******P*** < 0.001, *******P*** < 0.0001.

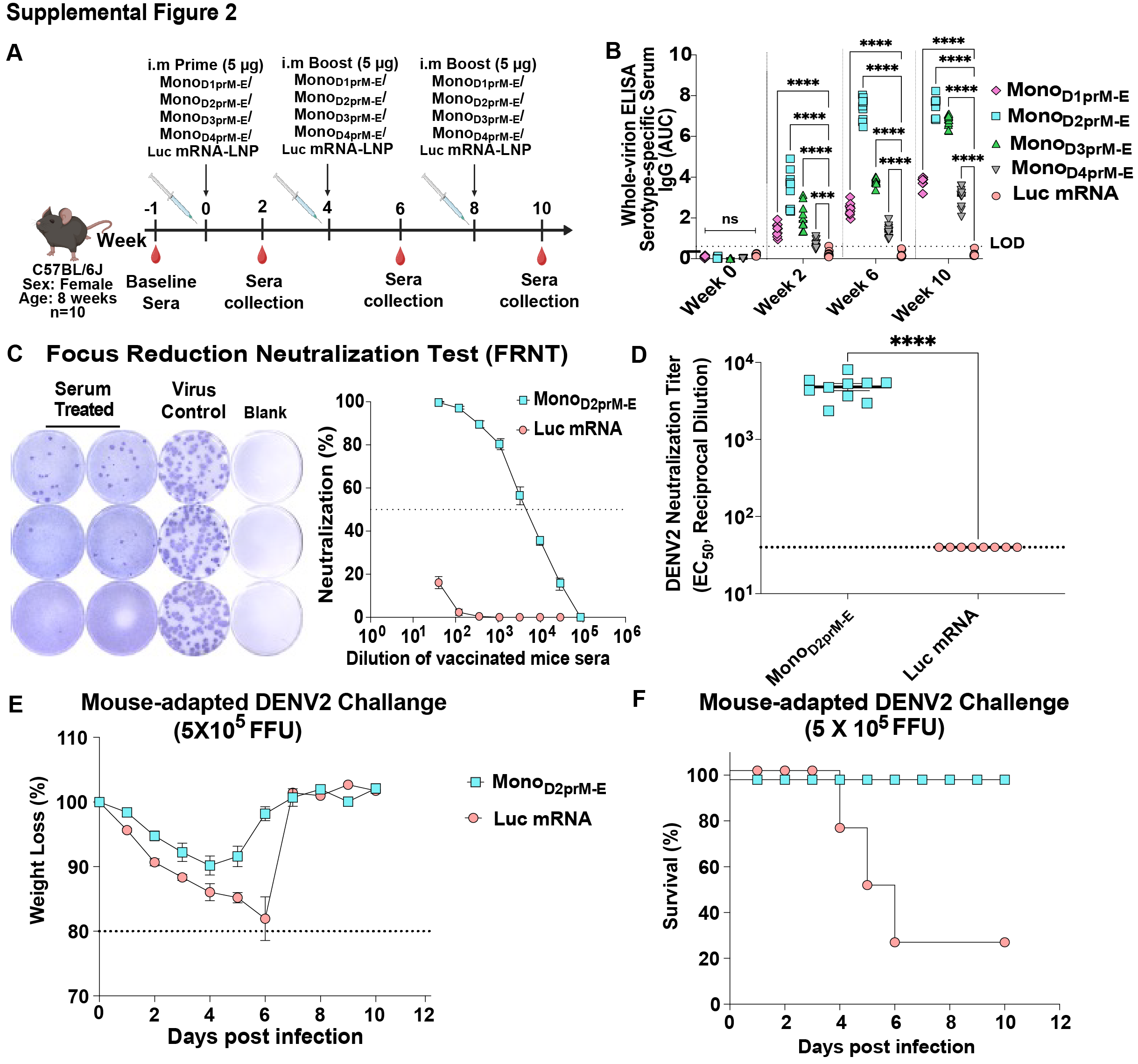

**Supplementary Figure 2:** **Immunogenicity and protective efficacy of DENV monovalent prM-E mRNA-LNP vaccination. (A)** Schematic of the immunization and serum collection schedule. Eight-week-old female C57BL/6J mice (n = 10 per group) were immunized intramuscularly (i.m.) with 5 μg of DENV1/ DENV2/ DENV3/ DENV4 monovalent prM-E (Mono_D1prM-E_/ Mono_D2prM-E_/ Mono_D3prM-E_/ Mono_D4prM-E_) mRNA-LNP or luciferase (luc) mRNA-LNP control at weeks 0, 4, and 8. Sera were collected at baseline and at weeks 2, 6, and 10. **(B)** DENV-specific whole-virion serum IgG responses measured by ELISA at the indicated time points and expressed as area under the curve (AUC). Sera from each monovalent vaccine group were tested against whole virions of the corresponding DENV serotype. The ELISA limit of detection (LOD) was defined as, LOD = mean of the blank + 3 × SD of the blank. **(C)** Representative focus reduction neutralization test (FRNT) images (left) and DENV2 neutralization curves (right) for sera from Mono_D2prM-E_- and luc mRNA-immunized mice. Neutralization is plotted as a function of serum dilution; the dotted horizontal line indicates 50% neutralization. **(D)** DENV2 neutralizing antibody titers (EC₅₀) in sera from Mono_D2prM-E_- and luc mRNA-immunized mice. Dashed lines indicate the assay LOD at a 1:40 dilution. Passive protection study using sera from mice immunized with Mono_D2prM-E_ or luc mRNA vaccines. Immune sera were transferred into type I interferon receptor knockout (IFNAR1) mice, which were challenged 18 h later with 5 × 10⁵ focus-forming units (FFU) of mouse-adapted S221 DENV2. **(E)** Body weight following DENV2 challenge, expressed as percentage of baseline body weight. The dotted horizontal line indicates 80% of baseline body weight. **(F)** Survival following DENV2 challenge. Comparisons between two groups were performed using a two-tailed Mann-Whitney U test. For analyses involving two factors with repeated measurements, statistical significance was assessed using a mixed-effects model (REML) with the Geisser-Greenhouse correction, followed by Tukey’s multiple-comparisons test. ns, not significant; ****P* < 0.001; *****P* < 0.0001.

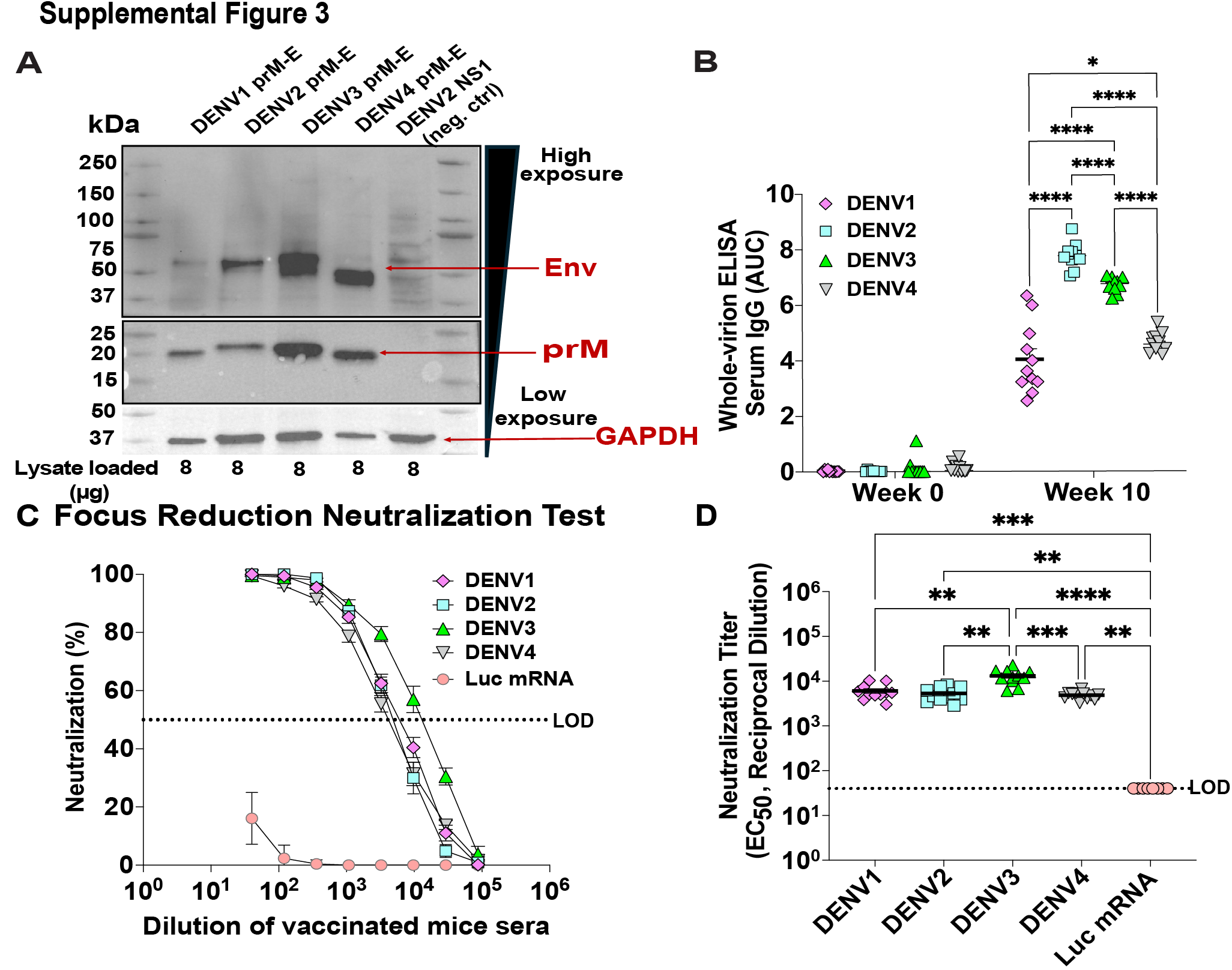

**Supplementary Figure 3: Tetravalent DENV PrM-E (Tet_prM-E_) mRNA-LNP vaccination induces binding and neutralizing antibody responses against all four DENV serotypes.** **(A)** Western blot analysis of cell lysates expressing the indicated DENV1, DENV2, DENV3, and DENV4 prM-E constructs, confirming expression of the precursor membrane (prM) and envelope (env) proteins. DENV2 NS1-expressing cell lysate was included as a negative control. GAPDH was used as a loading control. A total of 8 μg of cell lysate was loaded per lane. High and low exposures are indicated. **(B)** DENV-specific serum IgG responses following Tet_prM-E_ mRNA-LNP vaccination were assessed by whole-virion ELISA. Sera collected at baseline (week 0) and week 10 were evaluated for IgG binding to whole DENV1, DENV2, DENV3, and DENV4 virions. Antibody binding is expressed as area under the curve (AUC), with each symbol representing an individual mouse. **(C)** Focus reduction neutralization test (FRNT) showing neutralization of DENV1-4 by sera from Tet_prM-E_ mRNA-LNP-vaccinated mice. Neutralization is plotted as a function of serum dilution. Sera from luciferase (luc) mRNA-LNP immunized mice were included as a negative control. The dotted horizontal line indicates 50% neutralization. **(D)** DENV1-4 neutralizing antibody titers (EC₅₀) in sera from Tet_prM-E_ mRNA-LNP-vaccinated mice. The dotted horizontal line indicates the assay limit of detection (1:40). Each symbol represents an individual mouse. Comparisons among multiple groups were performed using the Kruskal-Wallis test followed by Dunn’s multiple-comparisons test. *P < 0.05, *****P*** < 0.01, ******P*** < 0.001, *******P*** < 0.0001.

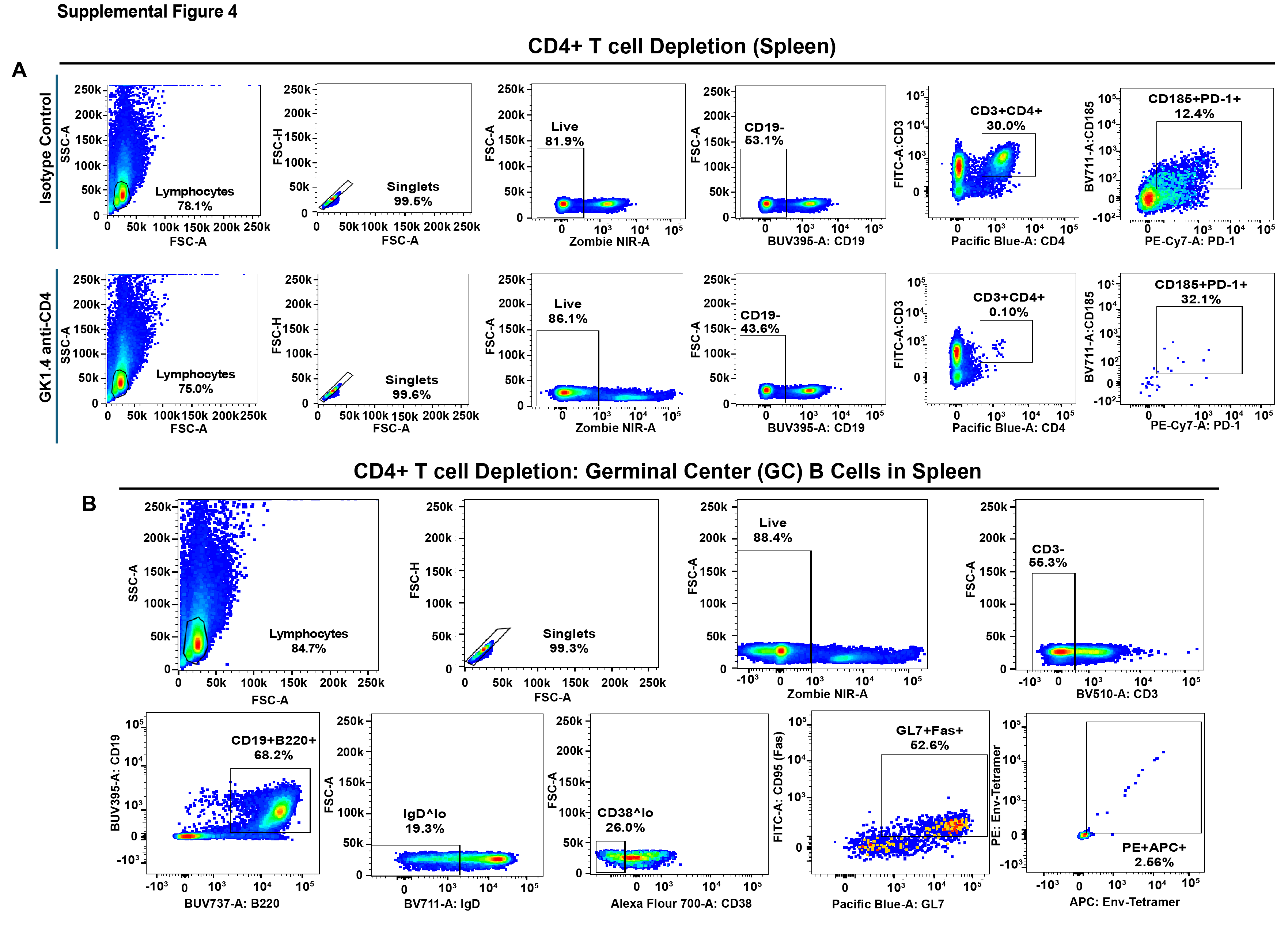

**Supplementary Figure 4: Flow cytometry gating strategy for assessment of CD4⁺ T cell depletion and germinal center (GC) B cell responses in the spleen.** Representative flow cytometry plots showing the sequential gating strategy for splenic immune cell populations from isotype control and GK1.5 anti-CD4-treated mice. **(A)** CD4⁺ T cell analysis. Lymphocytes were identified by forward- and side-scatter characteristics, followed by selection of singlets, live cells, and CD19⁻ cells. CD4⁺ T cells were subsequently identified based on CD3 and CD4 expression, followed by assessment of CD185 (CXCR5) and PD-1 expression. Representative plots demonstrate the marked reduction of the CD3⁺CD4⁺ population following GK1.5 treatment. **(B)** Germinal center (GC) B cell analysis. Following lymphocyte, singlet, and live-cell gating, CD3⁻ cells were selected and B cells were identified as CD19⁺B220⁺. Subsequent gates were applied to identify IgD^lo and CD38^lo populations, followed by assessment of the GC-associated markers GL7 and CD95 (Fas). DENV3 env-specific cells were subsequently identified within the indicated population. Numbers shown within gates indicate the percentage of cells within the respective parent population.

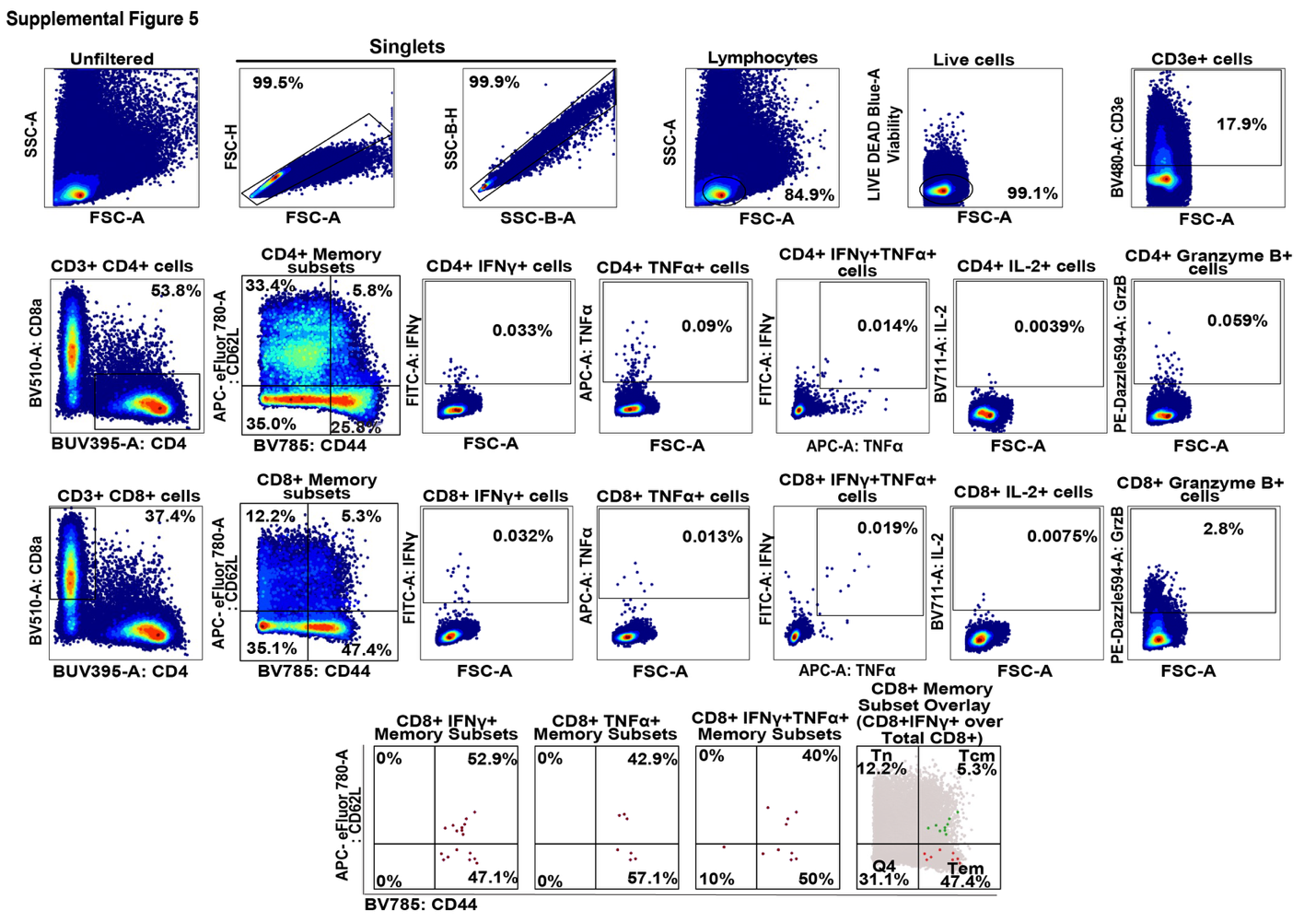

**Supplementary Figure 5: Flow cytometry gating strategy for characterization of T-cell responses in splenocytes following ex vivo stimulation.** Representative flow cytometry plots illustrating the gating strategy used to identify and characterize antigen-responsive CD4⁺ and CD8⁺ T-cell populations. Cells were sequentially gated to exclude doublets, followed by identification of lymphocytes, live cells, and CD3ε⁺ T cells. CD3ε⁺ T cells were subsequently separated into CD4⁺ and CD8⁺ T-cell populations. CD4⁺ and CD8⁺ T-cell memory subsets were defined based on CD44 and CD62L expression. Functional responses within CD4⁺ and CD8⁺ T cells were assessed by intracellular staining for IFN-γ, TNF-α, IL-2, and granzyme B, including identification of IFNγ⁺TNF-α⁺ double-positive cells. For responding CD8⁺ T cells, CD44 and CD62L expression was further evaluated within IFN-γ⁺, TNF-α⁺, and IFN-γ⁺TNF-α⁺ populations to characterize their memory phenotypes. Representative plots and gating boundaries are shown.

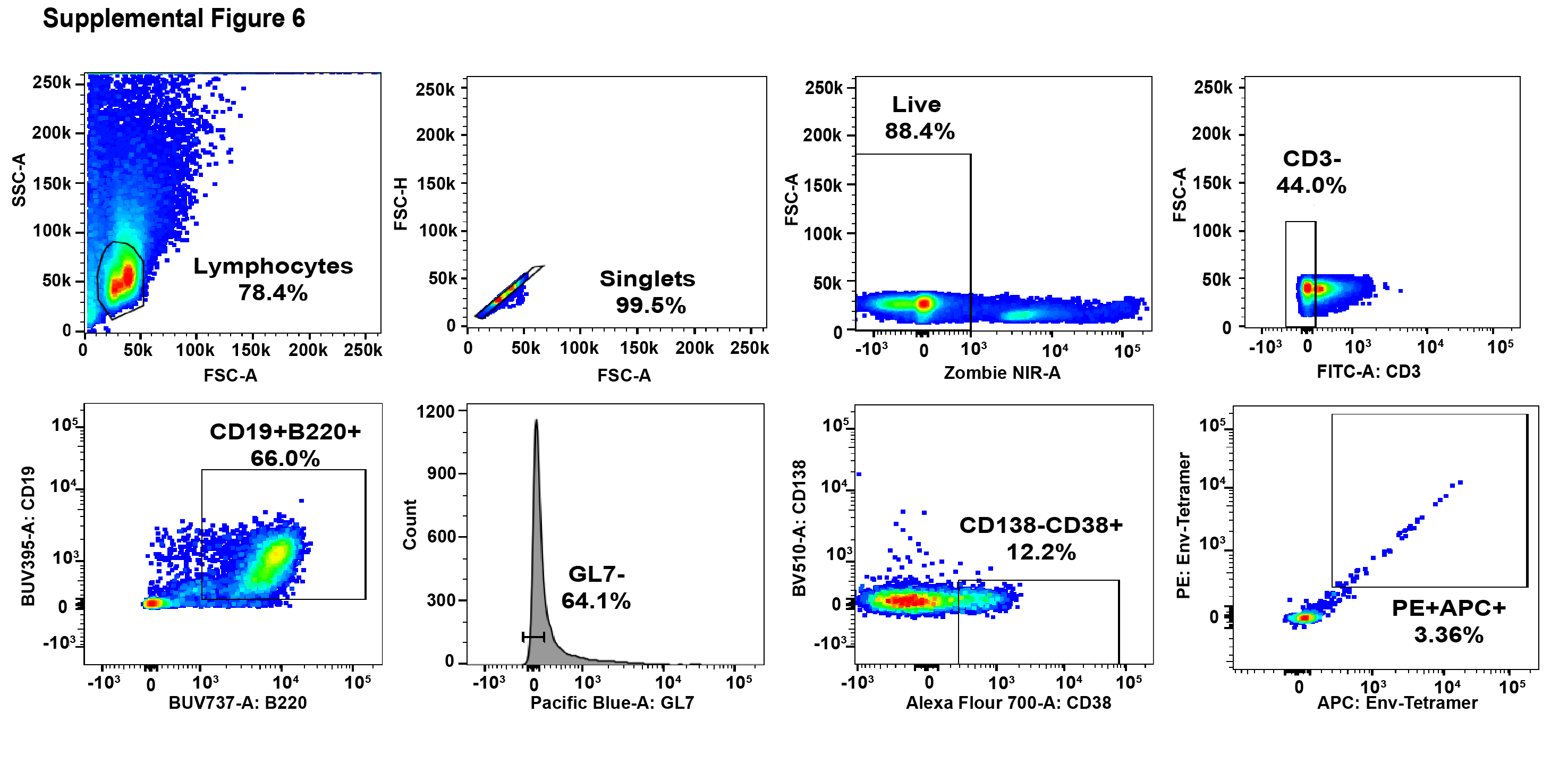

**Supplementary Figure 6: Flow cytometry gating strategy for identification of DENV3 env-specific memory B cells (MBCs) in the spleen.** Representative flow cytometry plots showing the sequential gating strategy used to identify DENV3 env-specific MBC populations in spleen collected at week 10. Lymphocytes were first identified based on forward- and side-scatter characteristics, followed by selection of singlets and live cells. CD3⁻ cells were then selected to exclude T cells, and B cells were identified as CD19⁺B220⁺. Subsequent gating was performed based on GL7, CD138, and CD38 expression. DENV3 env-specific MBCs were identified by dual binding to PE- and APC-conjugated env-tetramers. Numbers shown within gates indicate the percentage of cells within the respective parent population.

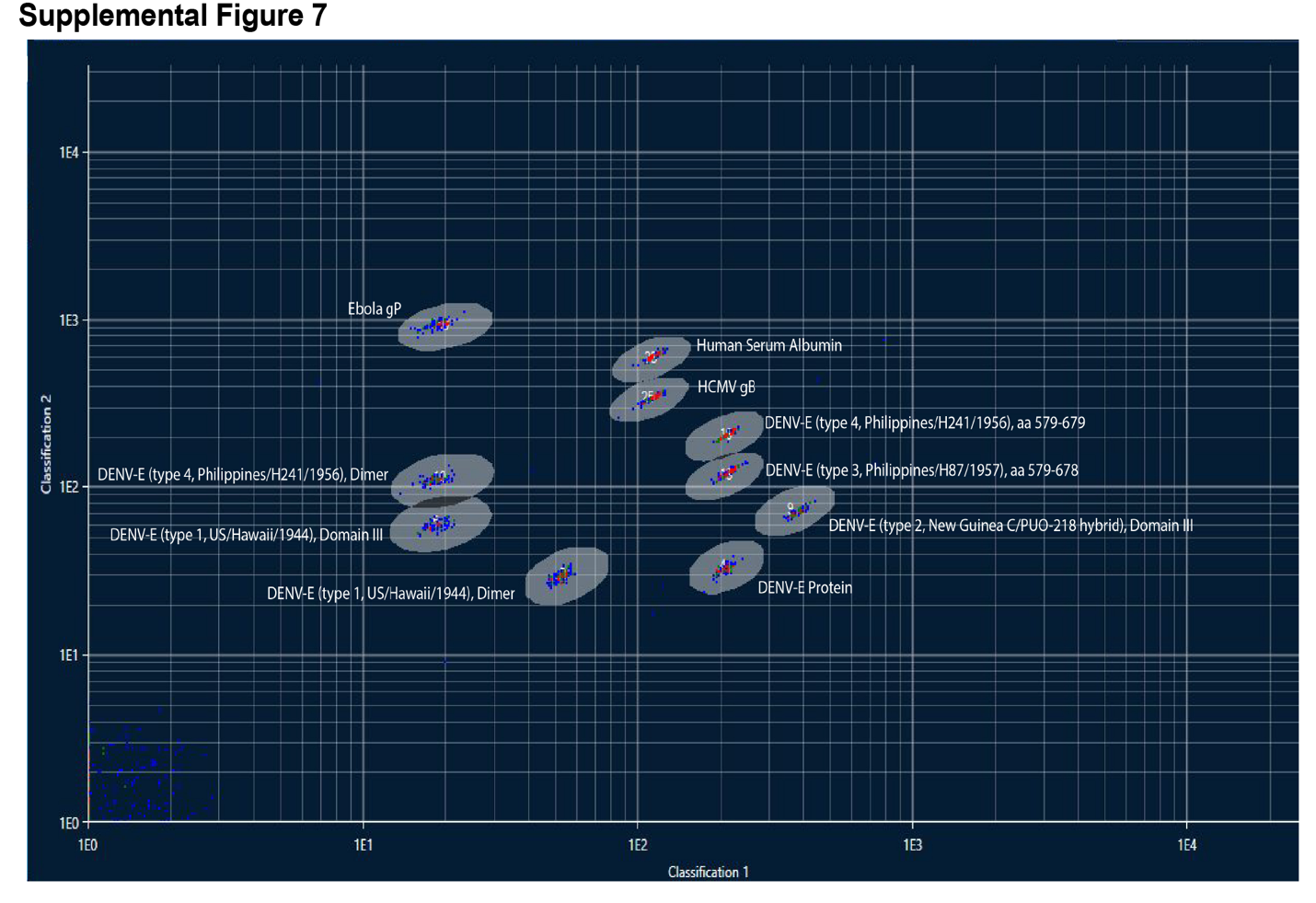

**Supplementary Figure 7: Antigen bead classification and gating for the multiplex systems serology assay.** Representative antigen-classification plot showing separation of the antigen-coupled bead populations used in the multiplex assay, including DENV antigens, HCMV gB, Ebola glycoprotein (gP), and human serum albumin.

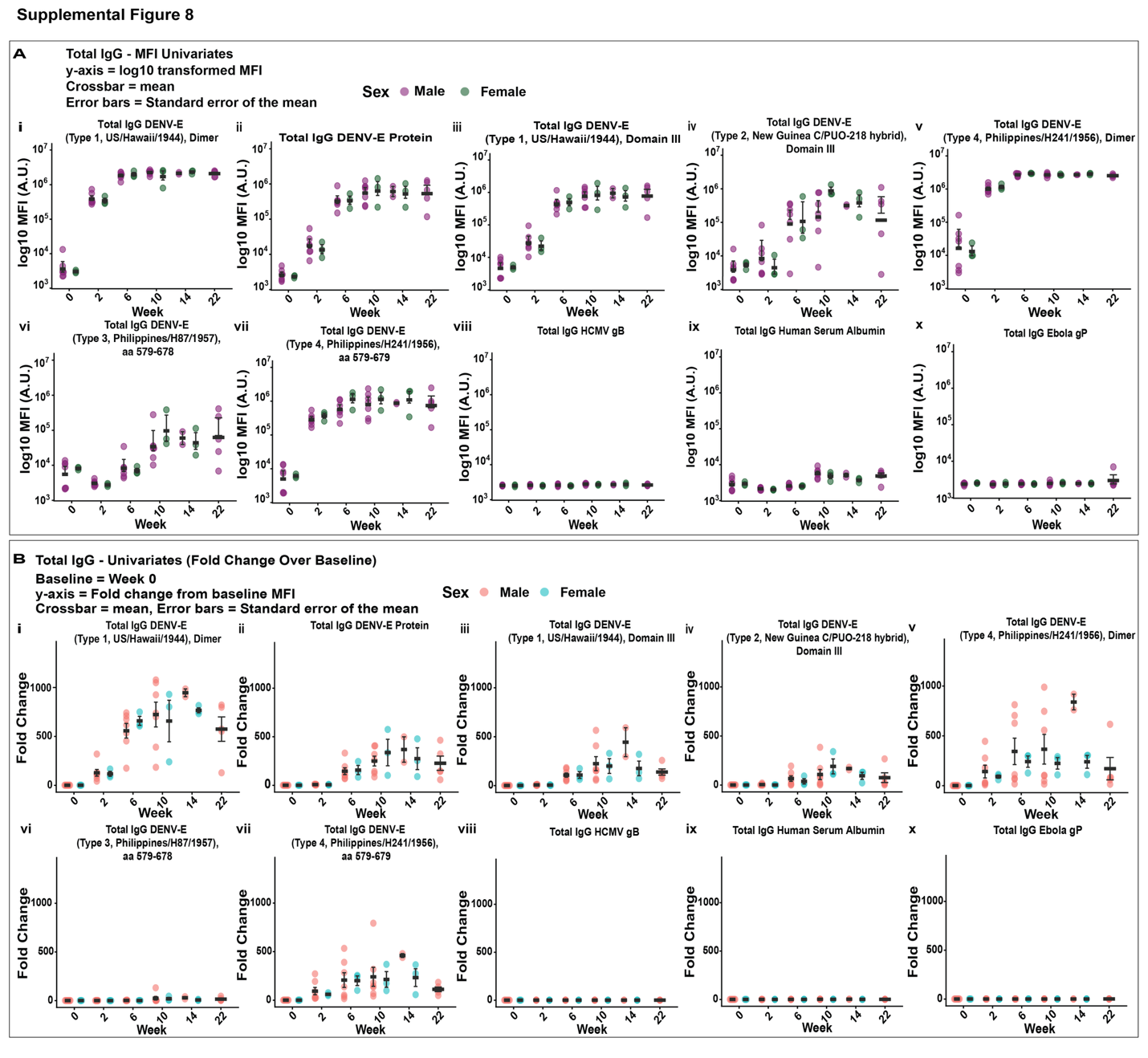

**Supplemental Figure 8: Longitudinal DENV-Specific IgG Responses. (A)** Longitudinal total IgG responses to individual DENV antigens and control proteins, expressed as MFI (arbitrary units, A.U.). **(B)** Corresponding total IgG responses expressed as fold change relative to baseline. In **(A)** and **(B)**, individual data points represent individual animals and are colored by sex; black symbols and error bars indicate the mean and standard error of the mean (SEM), respectively.

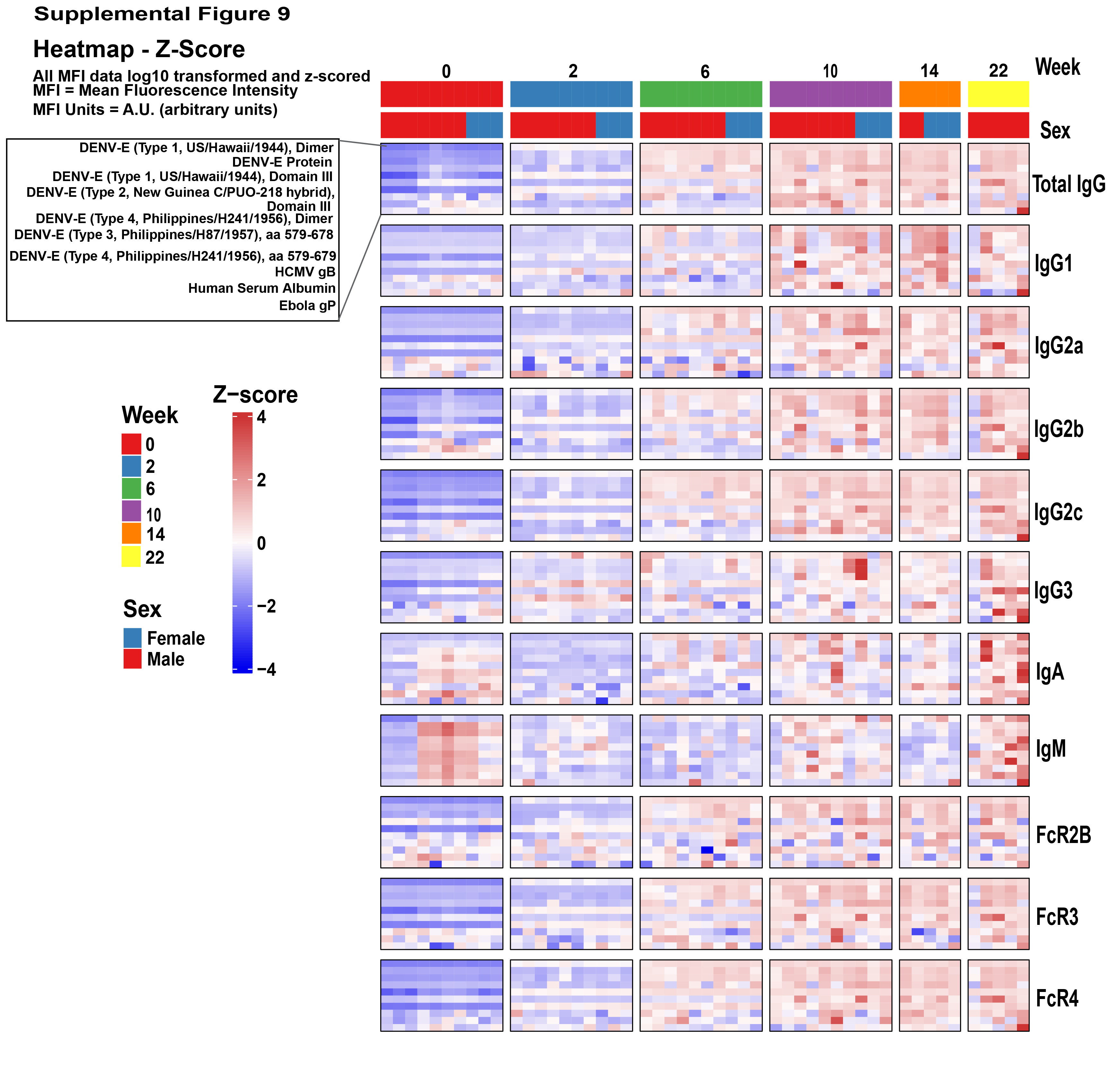

**Supplemental Figure 9: Sex-based differences in longitudinal systems serology profiles of DENV-specific antibody responses.** Heatmap of antibody and Fc receptor-binding profiles across study time points. Mean fluorescence intensity (MFI) values were log10-transformed and z-score normalized and are shown for Total IgG, IgG subclasses (IgG1, IgG2a, IgG2b, IgG2c, IgG3), IgA, IgM, FcR2B, FcR3, and FcR4. Columns represent individual samples, with annotations indicating week and sex, and rows represent the indicated antigens.

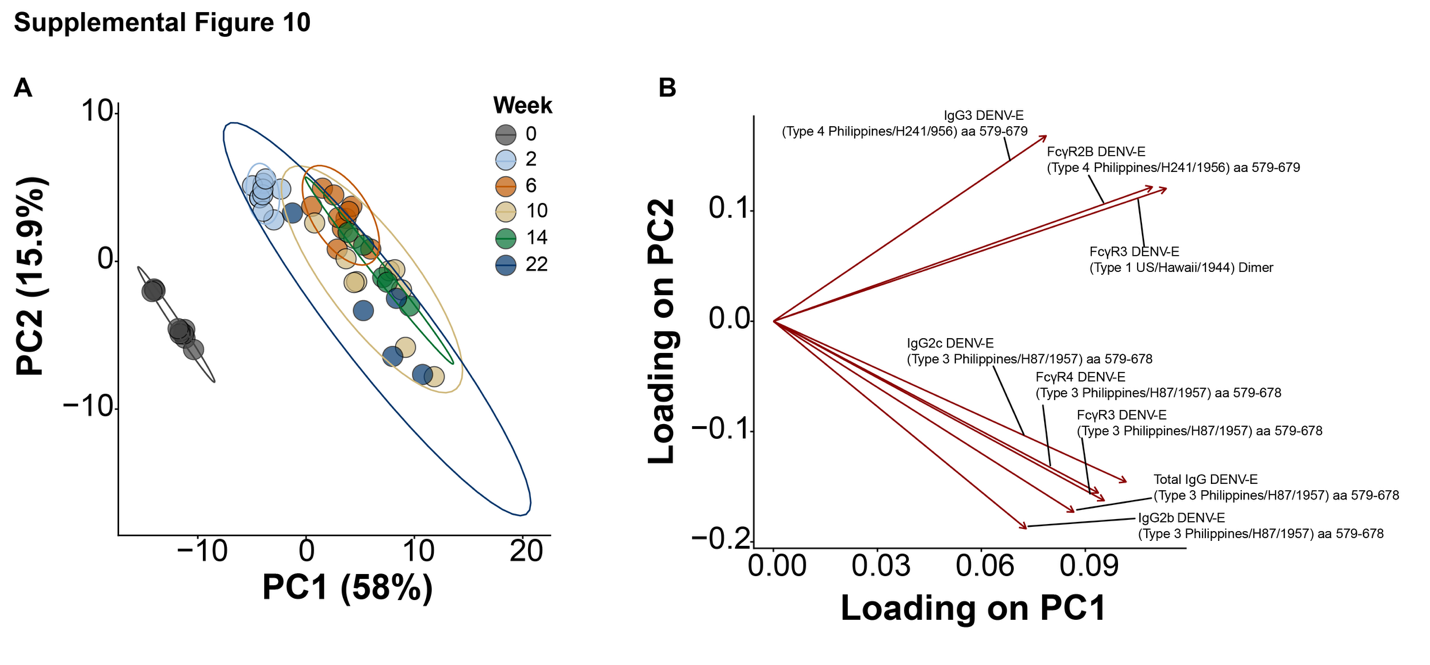

**Supplemental Figure 10: Principal component analysis of longitudinal systems serology profiles following tetravalent DENV prM-E (Tet_prM-E_) mRNA-LNP vaccination. (A)** Principal component analysis (PCA) of composite humoral profiles across longitudinal time points. PC1 accounted for 58% of the total variance and separated post-vaccination samples from baseline. **(B)** PCA biplot showing the top eight humoral features contributing to sample separation, including IgG subclass and Fcγ receptor-binding responses directed against DENV1, DENV3, and DENV4 envelope-derived antigens.
